# Analysis of RNA translation with a deep learning architecture provides new insight into translation control

**DOI:** 10.1101/2023.07.08.548206

**Authors:** Xiaojuan Fan, Tiangen Chang, Chuyun Chen, Markus Hafner, Zefeng Wang

## Abstract

Accurate annotation of coding regions in RNAs is essential for understanding gene translation. We developed a deep neural network to directly predict and analyze translation initiation and termination sites from RNA sequences. Trained with human transcripts, our model learned hidden rules of translation control and achieved a near perfect prediction of canonical translation sites across entire human transcriptome. Surprisingly, this model revealed a new role of codon usage in regulating translation termination, which was experimentally validated. We also identified thousands of new open reading frames in mRNAs or lncRNAs, some of which were confirmed experimentally. The model trained with human mRNAs achieved high prediction accuracy of canonical translation sites in all eukaryotes and good prediction in polycistronic transcripts from prokaryotes or RNA viruses, suggesting a high degree of conservation in translation control. Collectively, we present a general and efficient deep learning model for RNA translation, generating new insights into the complexity of translation regulation.

## Introduction

Computational analysis of protein-coding open reading frames (ORFs) in RNAs have traditionally relied on empirical rules such as sequence conservation pattern (1), the longest ORFs in one transcript (2) and initiation from the first AUG (3). However, these rules are over-simplified without considering the complex regulation by other factors, such as RNA structure and various regulatory *cis*-elements. As a result, previous efforts in *ab initio* prediction of translation sites achieved relatively low accuracy (4). In addition, RNA translation has recently been found in previously annotated non-coding RNAs (e.g., lncRNAs and circRNAs) (5–7), and can be initiated from alternative sites or non-canonical codons (8,9), resulting in non-annotated translation products that play critical and diverse cellular roles (6,8). Considering the complexities of translation control, the prediction of translation sites with high accuracy remains a challenging task, especially for poorly annotated genomes or in the context of complex regulatory networks. Rapid advances in artificial intelligence (AI) offer new hope for dissecting complex biological systems, such as alternative splicing (10) and protein folding (11), by uncovering hidden associations and unknown rules (12–15).

The complete synthesis of a protein requires accurate initiation and termination of translation by ribosomes at specific positions. In eukaryotes, translation initiation is generally mediated by the 5′-cap, which recruits various translation initiation factors and small ribosomal subunit to form preinitiation complex (16). This complex subsequently scans through the 5′-untranslated region (5′-UTR) to recognize the optimal start codon, which is determined by both codon identity and its immediate nucleotide context known as Kozak sequence GCCRCC<u>AUG</u>G (R represents purine) (3). However, in certain cases, the ribosome may skip some weak start codons, resulting in leaky scanning (3). In addition, a large fraction of genes contain several alternative translation initiation sites (TISs), but it is unclear how the ribosomes select one of these sites to initiate translation. Evaluating the strength of each TIS within one transcript and the general rules of selecting the authentic ORFs from possible decoys remain challenging. Therefore the systematic identification of translation sites (translation initiation and termination sites, or TIS/TTS) and the quantitative evaluation of their strength become a prerequisite for exploring the complexity of translation regulation.

New experimental techniques, including GTI-seq (global translation initiation sequencing) (17) and QTI-seq (quantitative translation initiation sequencing) (18), have been developed to identify TISs by directly capturing the stalled initiating ribosomes at single nucleotide resolution across the transcriptome. However, these methods were typically conducted in specific cell types, which limits the transcriptome-wide TIS identification. Several computational methods have also been developed to predict translation initiation sites from mRNA sequences. For example, a physicochemical-property based predictor using pseudo-trinucleotide composition was constructed to identify TIS in human genes (19), and a variety of machine learning methods have also been developed to predict the TIS by analyzing a short sequence fragment (e.g. 200nt) around the start codon of mRNA (20–23). However, such methods are biased because they are limited to the immediate sequence context surrounding the TISs. In other words, these approaches do not consider the entire non-coding regions of mRNAs (i.e. 5′-UTR and 3′-UTR) that play an important regulatory role in selecting the start codon (24,25), which in turn affects the accuracy of TIS predictions.

Here, we developed a deep learning method based on a multilevel dilated convolution network, named TranslationAI, to independently predict TISs and TTSs from full-length mRNA sequences alone. Leveraging the inherent structure of the genetic code, TranslationAI achieves high accuracy, surpassing a >99% Precision-Recall Area Under the Curve (PR-AUC) in predicting known TISs and TTSs in human transcriptome. Our model uncovered regulatory sequences in the UTRs that determine the strength of translation start and stop codons, and identified thousands of new ORFs in mRNAs or annotated lncRNAs. In addition, this model can be extended to other eukaryotes, prokaryote and several human viruses, suggesting a strong conservation in defining coding sequences across different domains of life. To facilitate further applications by other researchers, we developed a web tool (https://www.biosino.org/TranslationAI/) that is accessible to end users in predicting translation sites from RNA sequences. The application of TranslationAI extends beyond *de novo* prediction of TIS/TTS in any given transcripts, as it can also be used to evaluate the translation potential of a given RNA sequence.

## Results

### Modeling translation from full length mRNA with deep learning

We constructed a deep residual convolutional neural network, TranslationAI, using the full-length mRNA sequences as input to independently predict the TIS and TTS (Fig. 1A, see methods). Previous approaches only considered short sequence stretches around a potential start codon (20–22), or reported all potential ORFs in six frames with specified start codon (ATG), stop codon (TAG/TAA/TGA), and length cutoffs (ORFfinder in NCBI). In contrast, our deep learning model evaluates the potential of each position in a given mRNA to function as a TIS or TTS without using any prior knowledge of translation (e.g., the triplet or identity of genetic code). Such *ab initio* prediction enables the neural network to learn the hidden rules of translation from sequence context alone.

**Figure 1.**
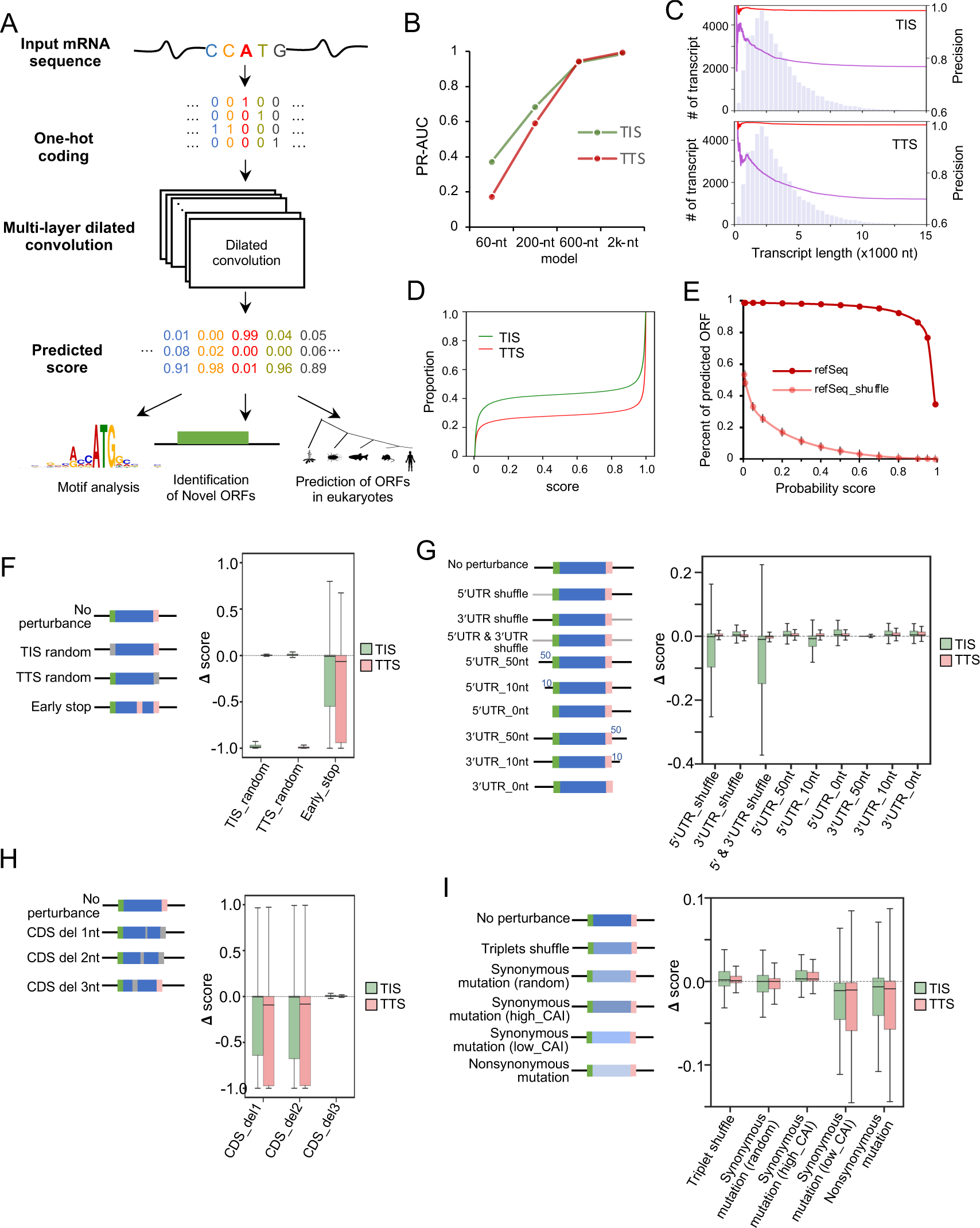
Construction of deep learning network for translation prediction. **A.** Flowchart of TranslationAI, a computational model for predicting translation initiation and termination sites with full length mRNA. For each position in the full length mRNA, TranslationAI-2k takes 60, 200, 600, and 2,000 nucleotides of flanking sequence as input, and predicts whether that position corresponds to a translation initiation site (TIS), translation termination site (TTS), or neither. **B.** Effect of input sequence context size on network accuracy. PR-AUC is the area under the precision-recall curve. The figure shows the performance of the network at four different input sequence context sizes. **C.** Relationship between transcript length and the positive rate of TIS/TTS for each transcript by comparing the number of positive TISs from TranslationAI-200 (purple line) or TranslationAI-2k (red line) to the total number of transcripts with lengths shorter than the given transcript. The distribution of transcript length is shown in the background as a histogram. **D.** Accumulative distribution of TIS/TTS score among all mRNAs. **E**. The number of ORFs predicted by TranslationAI-2k using mRNA and shuffled mRNA (mononucleotide shuffling) as input and different cutoff of TIS scores. The average number of ORFs predicted from multiple shuffling, along with error bars representing three times the standard deviation calculated from shuffled sequences. **F-I.** The features learned by TranslationAI. Systematic *in silico* perturbations on different regions of the mRNA, measured as changes (Δ score) in probability scores for the authentic TISs/TTSs. The perturbations include: **F**, replacement of TIS/TTS identity; **G**, changes in UTR length and sequence; **H**, frameshifts by deleting one, two, or three nucleotides; **I**, mutations in coding sequences. Blue box: ORF (open reading frame), black line: 5′-UTR and 3′-UTR, green line: TIS, pink line: TTS. Mononucleotide shuffling was performed for shuffling of 5′-UTR and 3′-UTR. Codon shuffling was performed for triplet shuffle, which maintained the amino acid composition. CAI: Codon Adaptation Index.

The deep residual neural network was constructed using the one-hot encoding of mRNA sequences as input (Fig. 1A). We built a 32-layer dilated convolution neural network architecture that produces an output matrix of the probability for the given position being a TIS, TTS or neither (Fig. S1, see methods). To obtain a comprehensive and well-curated dataset, we utilized the RefSeq gene annotation and extracted 47,098 protein-coding transcripts (47,098 TIS-TTS pairs). We trained the model using the transcripts on chromosomes 2, 4, 6, 8, 10-22, X, and Y, and tested the model with the transcripts from chromosomes 1, 3, 5, 7, and 9. The stringent top-k accuracy was used for TIS/TTS prediction. Specifically, the score cutoff was set at the point where the number of predicted sites is equal to the real number (i.e., the false positive rate equal to false negative rate, the score cutoff at this point is ∼0.5), which ensured that only the high confidence predicted sites were considered for downstream analysis.

To explore the effect of different input windows on test accuracy, we considered four input windows that include 60, 200, 600, and 2,000 (2k) nucleotides on both sides of a given position. Interestingly, the prediction accuracy improved dramatically with the increasing input window size, reaching >99% PR-AUC with a 2k nt input window on test dataset (Fig. 1B, Table S1). Considering that >70% of transcripts are longer than 2k nt (Fig. S2A), such high accuracy of TranslationAI prediction is particularly remarkable, suggesting that the identity of canonical TIS/TTS could be influenced by the sequences at thousands of nucleotides away, potentially through long-range interactions such as RNA structure or RNA binding proteins. Our findings further suggest that TranslationAI has learned to capture these interactions using sequence information alone.

To assess the impact of sequence context on the accuracy of TIS/TTS prediction, we compared the performance of the models trained on 200 and 2k nt sequence context. The 2k model achieved high prediction precision (>0.95) in canonical TIS/TTS prediction across a wide range of transcript lengths, however the precision of the 200 nt model decreases dramatically with increase of transcript length (Fig. 1C). This observation is in agreement with that earlier report that distal regions far away from the canonical TIS/TTS can affect the translation decision (24). With the 2k model, the score distribution showed a clear separation between the positive and negative predictions (Fig. 1D), suggesting that the model is robust under different cutoff selections. To further examine the false discovery rate (FDR), we used the 2k model to score potential TISs/TTSs in the shuffled mRNA sequences. At the probability score cutoff of 0.5 for both TIS and TTS, less than 5% of the ORFs in shuffled dataset passed the cutoff, yielding an empirical FDR < 0.05 (Fig. 1E).

### Key features learned by the AI model for accurate prediction

Encouraged by the near-perfect performance of our deep learning network in predicting canonical translation sites, we sought to examine the features learned by this “black box” model. We performed systematic *in silico* perturbations on different regions of mRNAs to measure their effects on reducing the probability scores of the authentic TISs/TTSs. Such perturbations can reflect how the AI model make the accurate translation prediction in a computational sense, however a different molecular mechanism may be used in cells to define translation initiation and termination sites as judged by experiments. Nevertheless, it would be informative to compare how a biological process like mRNA translation is perceived differently by the AI model and experimental tests.

We found that replacing annotated TISs or TTSs with random codons dramatically reduced the predicted scores of cognate sites (Fig. 1F, Table S2), suggesting this model had successfully learned the identity of start and stop codons from the sequence alone. Consistently, introducing a known stop codon into the coding sequences (CDS) led to a drastic reduction in predicted scores of both TISs and TTSs. In addition, shuffling the 5′-UTRs significantly affected the corresponding TIS prediction but not TTS, whereas shuffling the 3′-UTR had little effect on TIS/TTS prediction (Fig. 1G, Table S2). Notably, when both 5′-UTR and 3′-UTR were simultaneously shuffled, the TIS prediction score was lower compared to the cases when only the 5′-UTR was perturbed (Fig. 1G, Table S2), underscoring a potential synergistic role of 5′-UTR and 3′-UTR in TIS selection. Curiously, the deletions of 5′- or 3′-UTRs had less effect on TISs/TTSs prediction by this model, with near perfect prediction for transcripts with only 50 nt left in the UTRs (Fig. 1G, Table S2), however the underlying reason is unclear. Finally, random deletions of one or two nucleotides within CDS dramatically reduced the prediction scores, whereas random deletion of three nucleotides in CDS had little effect (Fig. 1H, Table S2), suggesting that the model was capable of learning the triplet rule of the genetic code by itself.

To further examine the impact of codon usage on our predictions, we introduced a series of perturbations to the genetic codons of CDS. First, maintaining the same amino acid compositions, the in-frame triplet shuffling or random synonymous substitutions had little effect on the prediction (Fig. 1I, Table S2). We further replaced codons with synonymous counterparts possessing either higher or lower Codon Adaptation Index (CAI). Notably, higher CAI substitutions increased prediction scores for both TIS and TTS, while lower CAI substitutions decreased these scores. These findings indicate that the AI model utilizes codon choice as a crucial factor in making predictions. Furthermore, we observed that nonsynonymous substitutions caused a small but statistically significant reduction in prediction scores. Collectively, these results suggest that the TranslationAI network has learned the intricate rules of RNA translation from a subset of the transcriptome without any prior knowledge.

To further investigate how other sites in the full-length mRNA contributed to the TranslationAI prediction, we calculated the feature importance of different positions of specific transcript in predicting TIS and TTS using an occlusion sensitivity analysis. Our results reveal that, while the sequences at the site of TIS and TTS contribute strongly to the prediction, the other positions at different regions also have various contributions (Fig. S2B). Remarkably, the importance of nucleotide at different positions relative to the TIS/TTS is dependent on the specific sequence contexts (i.e., the positional feature importance is different for each specific mRNA). For instance, in the case of a short mRNA GPX6, we observed a great impact of TIS identify on TTS prediction. In another case of a long mRNA PPP6R2, we found that a position in the 5¢-UTR shows high contribution to the TIS prediction. More specifically, when the original sequence “TAA” is masked into “NAA” or mutated, the score of the original TIS was greatly decreased (Fig. S2B).

In the RefSeq dataset, ∼100% of the annotated ORFs follow the triplet codon rule (length=3N), 98% are the longest ORFs in the transcripts, and ∼40% use the first AUG triplet as their start codon, raising the possibility that only using these simple rules may be sufficient to make prediction. To determine the extent to which TranslationAI relies on these simple rules, we generated *in silico* permutations in the test dataset and measured their effects on prediction precision. The introduction of frameshift permutations significantly reduced the prediction precision (Fig. S2C), however >50% of transcripts were still predicted to use original TIS/TTS despite breaking the 3N rule (i.e. ORF length ≠ 3N), suggesting that the rule of triplet codon is important but not mandatory for TIS/TTS prediction by TranslationAI. In addition, when ORF length was reduced by introducing premature stop codons or frameshifts, TranslationAI predicted the longest ORF in ∼60% cases (Fig. S2D), again suggesting that such a simple rule was partially followed by this model. Finally, the preference for the first AUG was also affected by several permutations throughout the transcript, including changes in TIS and the introduction of early stop or frameshift (Fig. S2E). Interestingly, shortening the length of the 5′-UTR increased the ratio of first AUG usage, whereas shuffling of the 5′-UTR reduced it, suggesting that the 5′-UTR plays a critical role in TIS selection (24,25). Collectively, these results indicate that TranslationAI has learned additional features beyond simple traditional rules (i.e., the longest ORF, triplet code, and preference of the first AUG) for accurate prediction, implying that the adjacent sequences near TIS/TTS also play crucial roles in selecting translation start and stop sites.

We found no correlation (Spearman’s correlation coefficient close to zero) between the predicted score of TIS/TTS and the number of transcript isoforms containing the same TIS/TTS sequences (Fig. S2F), suggesting that the scores of this model were not biased by the training data. Additionally, the TIS and TTS scores in the same ORF had a weak but statistically significant positive correlation (Spearman’s correlation coefficient r = 0.2, p < 1*10^-16^) (Fig. S2G), although they were predicted independently. Interestingly, the TIS/TTS scores by TranslationAI showed a positive correlation with the translation efficiency estimated by the ratio of read density from Ribosome profiling (Ribo-seq) normalized by mRNA abundance (8) (Fig. S2H and S2I). This consistency aligns with previous observations (Fig. 1I), where synonymous substitutions in CDS with higher CAI exhibits higher prediction scores, while substitutions with lower CAI shows lower prediction scores. This correlation implies that that these predicted scores partially reflect their endogenous translational activities.

### TranslationAI scores reveal new rule of codon selection for translation termination

We next sought to examine the hidden information of TranslationAI scores by comparing the key features between the strong and weak annotated TIS/TTS sites (the top *vs.* the bottom 5th percentile). Interestingly, the sequences surrounding strong TISs/TTSs are more conserved than those around weak TISs/TTSs (Fig. 2A), implying additional selective pressures for translation. Additionally, we examined the ribosome occupancy on the mRNAs with different TIS/TTS scores using Ribo-seq data from iPSC and iPSC-derived cardiomyocytes (8,26). Our analysis revealed that the transcripts with strong TISs or TTSs exhibit a higher ribosome peak at the start or stop positions (Fig. 2B-2C, Fig. S3A-S3B), which is consistent with their higher activity in mediating translation initiation or termination. These results suggest that the predicted scores of TISs/TTSs reflect their activities in regulating translational initiation/termination. In addition, the transcripts with weak TIS/TTS generally have a longer 5′-UTR than those with strong TIS/TTS (Fig. S3C), but such length difference was observed in the 3′-UTRs only for TTS with different strengths. However the implication behind such length difference is unclear.

**Figure 2.**
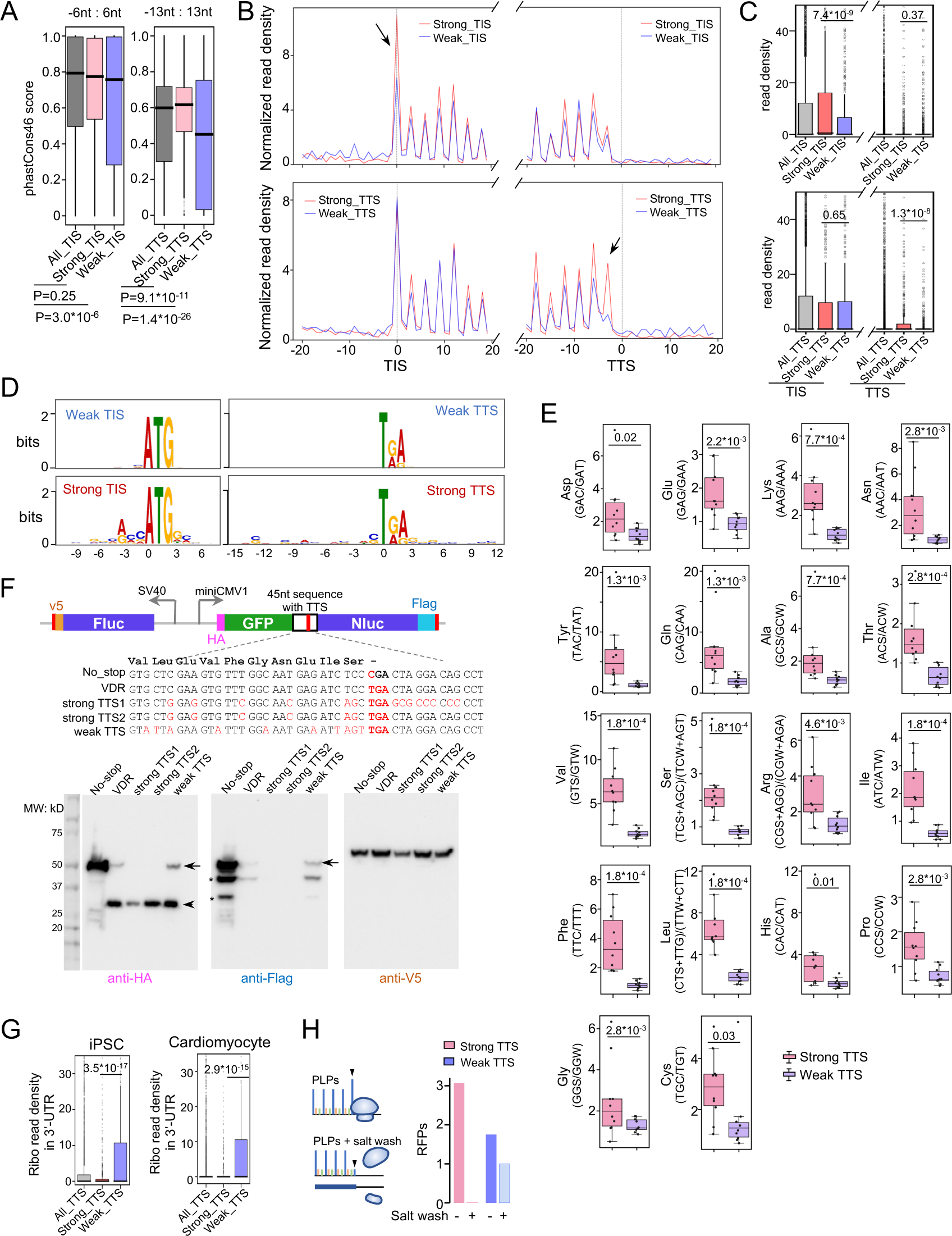
Evaluation of predictive features in TranslationAI. **A.** Conservation analysis of all TIS/TTS, strong TIS/TTS, and weak TIS/TTS regions. P-values were calculated using Mann–Whitney U test. **B.** Metagene analysis and **C.** boxplot of reads density on transcripts containing strong (red) and weak (blue) TIS/TTS (-20nt : +20nt) in iPSC cell line. P-values were calculated using Mann–Whitney U test. **D.** Motif analysis of strong and weak TIS/TTS. **E.** The codon distribution at the -30nt position of stop codon. The ratios of the same amino acid with C/G and A/T at the third position of each codon before strong and weak stop codons was quantified. P-values were calculated using standard T-test. **F.** Validation of motifs around strong TTS. Known readthrough stop codon from VDR and their mutations (without change of amino acids) were tested for translation readthrough. No_stop: stop codon of VDR (TGA) was mutated to CGA; strong TTS1: stop codon of VDR was mutated to a strong TTS by changing the upstream 27nt and downstream 12nt sequences; strong TTS2: stop codon of VDR was mutated to a strong TTS by changing the upstream 27nt sequence; weak TTS: stop codon of VDR was mutated to a weak TTS by changing the upstream 27nt. See supplementary information in Table S3. **G.** Boxplot analysis of reads density on 3′-UTRs from transcripts with strong/weak TTS in iPSC cell line and iPSC-induced Cardiomyocyte cell line, respectively. **H.** Normalized read density (by abundance of the transcript) at the last P-site before stop codon of in vitro platelet-like particles (PLP) and PLP treated with high salt.

We further examined the enriched sequence motifs around the strong *vs.* weak TISs/TTSs (from -30 nt to +33 nt) (Fig. 2D). As expected, we found an enriched motif RCC<u>ATG</u>GC (R = A or G, with start codon underlined) at the strong TISs, resembling the Kozak sequence that typically enhances translation initiation (27,28). We also found a novel motif S<u>TGA</u>G at the strong TTSs but not the weak TTSs (S = C or G, with stop codon underlined, Fig. 2D). Surprisingly, the fragments surrounding the strong TTS are generally biased towards CG-rich sequences, implying that the C/G-rich region may mediate more efficient translation termination. Moreover, the C/G enrichment showed a triplet periodicity before the strong TTSs (Fig. 2D), with the third position of a codon being most C/G biased. Such codon bias probably reflects the evolutionary selection under the constrain of protein sequences, suggesting an unappreciated role of codon usage in translation termination.

To confirm this codon bias, we analyzed the 30-nt region immediately upstream of the stop codon of human mRNA (equivalent of 10 codons). Consistent with the motif enrichment, the synonymous codons with C/G at the third position were preferably used at the upstream of strong TTSs (Fig. 2E, Fig.S3D). Remarkably, such codon bias was statistically significant for all 18 amino acids with multiple codons (Fig. 2E), particularly for the amino acids with 4-6 codons (e.g., Thr, Val and Ser etc., Fig. S3D). Together, these results reveal an intriguing role of codon selection in translation termination, and indicate that the TranslationAI has learned this codon bias to make a TTS prediction.

To experimentally validate the unexpected role of codon bias in translation termination, we designed a series of translation readthrough reporters with different TTSs (Fig. 2F). This reporter contains two independent transcription units, one encoding the firefly luciferase (Fluc) as a transfection control, and the other encoding a fusion protein of GFP and Nano-luciferase (Nluc) separated by a set of short variable sequences (45 nt) containing candidate TTSs of different strengths (Fig. 2F). As a control, we inserted the sequence around the terminal codon of the human vitamin D receptor (VDR) gene, which is known to undergo translation readthrough (29). We further introduced different synonymous mutations before the termination codon of VDR to either strengthen or weaken the TTS, and assayed the levels of mRNA and translation products from these reporters (Fig. 2F and S3E). The results showed that introducing the C/G-rich synonymous mutations in VDR (i.e., generating a strong TTS) eliminated the product from translation readthrough (indicated by arrows, Fig. 2F). Conversely, a weak TTS mutation with A/U-rich synonymous codons before the stop codon increased the translation readthrough. This experiment directly supported our finding on the role of codon selection in translation termination. As an independent validation, we re-analyzed the Ribo-seq data from two distinct cell lines to examine potential translation read through (8).

As expected, the ribosomal density in the 3′-UTRs following weak TTSs was significantly higher than that following strong TTSs (Fig. 2G), suggesting a translation “leakage” following the weak termination sites. To validate this observation, we conducted an analysis of the ribosome profiling data, in which the cells were subjected to a high-salt wash to release vacant ribosomes lacking a nascent polypeptide(30). The results demonstrated a distinct pattern: the high-salt wash effectively released the ribosome signal at the last P-site of strong TTS, while no such release was observed at weak TTS (Fig. 2H). This specific response is consistent with above findings that ribosomes associated with weak TTS may undergo translation readthrough, allowing continued translation beyond the canonical termination site, whereas ribosomes at strong TTS are more likely to be released, terminating translation at the expected site.

### Alternative TIS predicted by TranslationAI

Certain mRNAs may use alternative TIS to produce additional protein isoforms with extended N terminals (31), therefore we further examined if the TransaltionAI can predict alternative TISs that may compete with the annotated sites (see methods). Based on the distribution of TranslationAI scores (Fig. 1D), we used the score cutoff at 0.1 to increase the sensitivity of the alternative TISs prediction and identified 5336 alternative TISs in total. Interestingly, the numbers of alternative TIS identified by TranslationAI is negatively correlated with the strength of the annotated TISs in the same transcript (Fig. S3F), suggesting that alternative translation initiation may happen in the mRNA without dominant TIS. Specifically, ∼55% of mRNAs with weak annotated TISs contain an alternative TIS (1340 out of 2416 mRNAs), whereas the alternative TISs were found in less than 3% of mRNAs with strong annotated TISs (68 out of 2352 mRNAs). In addition, the score difference between the annotated TISs and the alternative TIS was much smaller in the mRNAs with weak TISs (Fig. S3G, left), whereas the mRNAs with strong annotated TISs generally lack alternative TISs with a competitive score (Fig. S3G, right). A consistent result was also found when we only considered the alternative TISs in the same reading frame with the annotated TISs (Fig. S3G). These results collectively suggested that the mRNAs with weak annotated TISs may also be translated from an alternative start site.

Further analysis suggested that the sequences at predicted alternative TISs are less conserved than the annotated TISs (Fig. S3H), suggesting that these alternative TISs may be emerged more recently during evolution. The predictions of alternative TISs were further supported by the Ribo-seq data obtained in two different cell lines (8), where the ribosome density in the 5′-UTRs could reflect the noncanonical translation initiation driven by alternative TIS. Consistently, the transcripts with a predicted alternative TIS showed significantly higher ribosomal read density in their 5′-UTRs compared to the mRNAs without alternative TIS (Fig. S3I), suggesting a leaky translation in the UTRs due to alternative translation initiation.

### TranslationAI identifies non-canonical ORFs in human transcriptome

It was recently reported that non-coding RNAs in the human transcriptome or the non-coding regions of mRNAs may contain unannotated ORFs that are translated into new proteins or small peptides (5,8). Therefore, we next used TranslationAI to examine the noncanonical ORFs in human transcriptome. Using the stringent cutoff for positive TISs/TTSs (i.e., same cutoff in mRNAs), we were able to identify 4620 noncanonical ORFs defined by new TIS-TTS pairs (Fig. 3A). These newly predicted ORFs included 673 upstream ORFs (uORFs, exemplified in Fig. 3B), 127 downstream ORFs (dORFs, exemplified in Fig. 3B), 26 overlapping ORFs with different reading frames from the annotated ORFs (Table S4). To validate the predicted non-canonical ORFs, we compared our prediction with uORFdb that is constructed from comprehensive and meticulous curation of uORF-related literature (32). The analysis revealed that a remarkable 569 out of 673 (85%) uTISs were also corroborated by uORFdb. In addition, 30 out of 673 (4%) uTISs were experimentally supported in TISdb using ribo-seq data from HEK293 (33) (Table S4).

**Figure 3.**
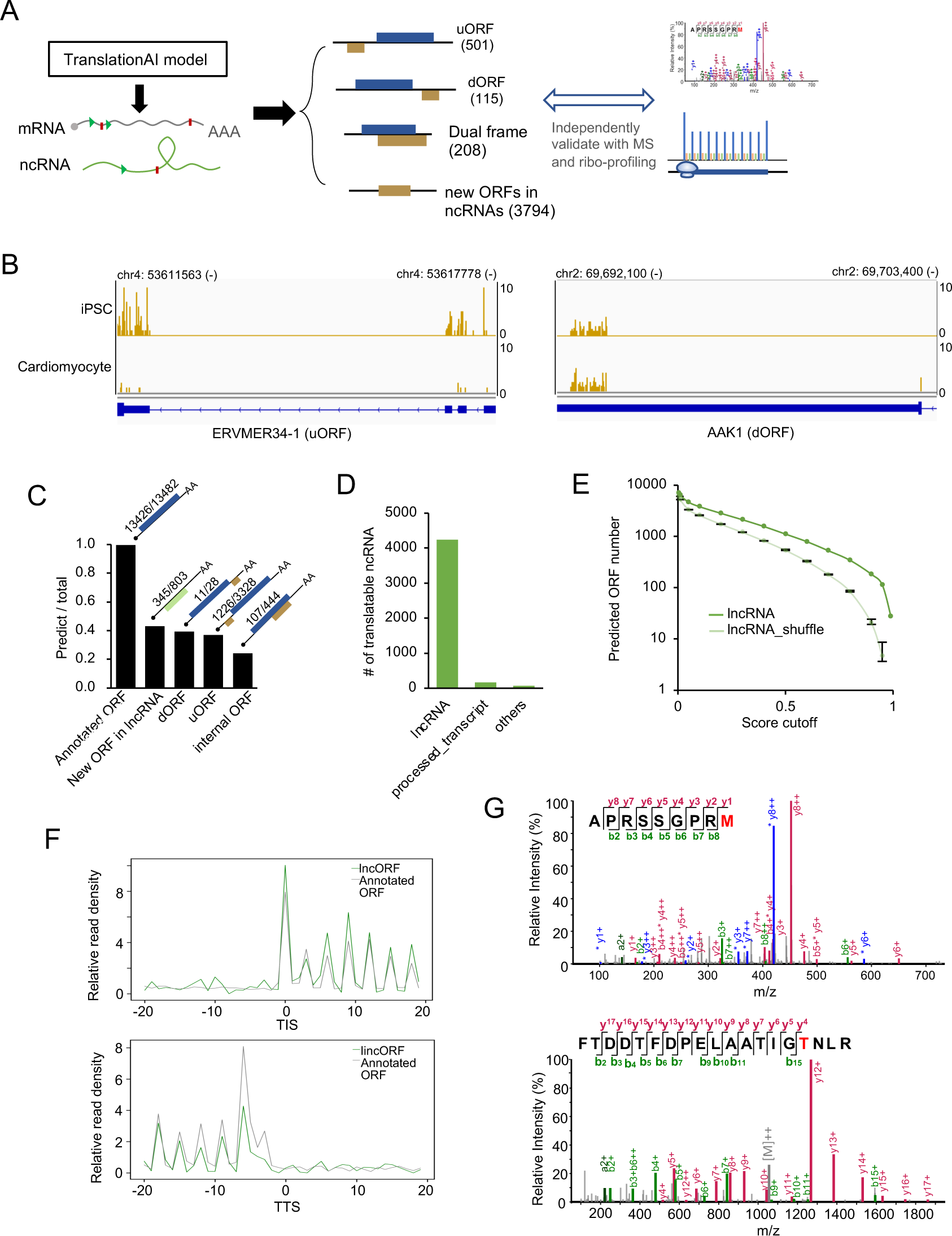
Identification of non-canonical ORFs in human transcriptome. **A.** Flowchart for identifying non-canonical ORFs, including upstream ORFs, downstream ORFs, dual coding ORFs, and new ORFs from non-coding RNAs. **B.** Example ribosome footprints of a uORF from ERVMER-1 and dORF from AAK1. **C.** The ratio of various types of predicted TISs identified by another published dataset derived from ribosome profiling assays. **D.** The number of newly identified translatable ncRNA in annotated lncRNAs, processed transcripts and other transcripts without known ORFs. **E.** The number of ORFs predicted by TranslationAI-2k using non-coding RNA and shuffled non-coding RNA sequences as input. The average numbers of ORFs predicted from lncRNAs and shuffled lncRNAs (mononucleotide shuffling, as control) were shown, with error bars representing 3×standard deviation calculated from shuffled sequences. **F.** Metagene analysis of translatable lncRNAs (green line) and control (grey line, mRNAs with the same ORF length distribution of predicted ORFs from lncRNAs). **G.** The MS/MS spectra of peptides from two ncRNAs: lncRNA ENST00000609975 (APRSSGPRM) and antisense RNA ENST00000413405 (FTDDTFDPELAATIGTNLR). The annotated b- and y-ions are marked in red and green, respectively.

The model also identified 3794 new ORFs from annotated noncoding RNAs (Fig. 3A, Table S5). These results demonstrate the potential for deep learning approaches to uncover previously unrecognized translation events and shed light on new regulatory mechanisms in gene expression. We next validated the prediction by comparing the predicted TISs in new ORFs with all TISs experimentally identified *via* ribosome profiling (8). We found that TranslationAI accurately predicted 99.6% of the annotated ORFs, however a smaller fraction of the noncanonical ORFs supported by Ribo-seq were successfully predicted by TranslationAI (39% of uORFs, 37% of dORFs, 24% of dual reading frame, and 43% new ORFs in lncRNAs were identified, Fig. 3C), suggesting that this model is biased towards canonical TISs/TTSs. Since the training set only contain canonical TIS/TTS information, the TranslationAI model may have learned limited features of non-canonical TISs/TTSs (see discussion).

We further focused on the thousands of predicted new ORFs from non-coding RNAs (ncRNAs), which are annotated in ENSEMBL as lncRNA, antisense RNA, miRNA precursors, processed transcripts, etc. (Fig. 3D). The shuffled sequences of all lncRNAs were analyzed using the same deep learning model as a background control. We found that the TranslationAI indeed predicted much more ORFs in the “non-coding” RNAs than their shuffled counterparts (Fig. 3E). While previous reports suggested that some lncRNAs can be translated into small peptides (5–8), detailed analysis of these noncanonical ORFs was inadequate. Our results showed that the newly predicted ORFs in lncRNA are significantly longer (Fig. S4A) and more conserved in 46 vertebrates (Fig. S4B) compared to the control ORFs defined by the longest fragments between ATG and TAA/TAG/TGA in all lncRNAs, suggesting a potential functionality. However, the TISs/TTSs from translatable lncRNAs have lower scores and are less conserved compared to canonical TISs/TTSs from mRNAs (Fig. S4C), implying that the translation efficiency of these newly identified ORFs may be lower than those from annotated mRNAs. This could partially explain why these ‘lncRNAs’ were originally defined as non-coding RNAs.

To validate the prediction of these new ORFs, we re-analyzed the Ribo-seq data from iPSC and cardiomyocytes (8). In total 64 newly predicted ORFs by TranslationAI were supported by Ribo-seq signals in either cell line (Table S5). The ribosome density of translatable lncRNAs closely resembled the footprints from control mRNAs, with strong trinucleotide periodicity that is indictive of active translation (Fig. 3F and Fig. S4D). In addition, mass spectrometry (MS)-based proteomics confirmed the stable expression of 191 non-canonical peptides (Table S5, with two examples shown in Fig. 3G). Among them, the ORFs in 10 lncRNAs were supported by both Ribo-seq and MS. The low number of experimentally validated ORFs may be contributed by several factors, including the cell line specific expression of most lncRNAs, the generally low expression level of lncRNAs, and the limited recover rate in both Ribo-seq and MS experiments.

Notably, we have correctly predicted 10 out of the 13 translated lncRNAs validated by independent functional studies, including the HOXB-AS3 that was reported to encode a 53aa peptide (34), the LINC00961 that encodes a 90aa peptide involved in the mTOR activation (6), and the RP11-132A1.4 that encodes a functional peptide of 124aa (8). All of these three lncRNAs were mis-annotated in RefSeq, probably because their expression and translation are limited to specific cell types. It is also worth noted that the predicted ORFs in annotated lncRNAs generally have low scores in our model (Table S5), which could be attributed to the short ORF length or the presence of a long 5′-UTR. Nevertheless, these results highlight the potential application of AI model in the future study of translatable lncRNA.

### TranslationAI accurately predicts TISs of other eukaryotes and viruses

An “universal genetic code” is used from bacteria to mammals, suggesting that translation machinery is highly conserved across different organisms. Therefore, we sought to examine if the model trained on human transcriptome can also predict canonical translation sites in other organisms. The accuracies of translation predictions for all eukaryotes tested (Human, Mouse, Zebrafish, Drosophila, Arabidopsis and budding yeast *S. cerevisiae*) were remarkably high (> 90% for all predictions, except that the yeast TIS prediction accuracy is 89%, Fig. 4A), suggesting that our model has learned the essential features generally required for translation initiation and termination in all eukaryotes.

**Figure 4.**
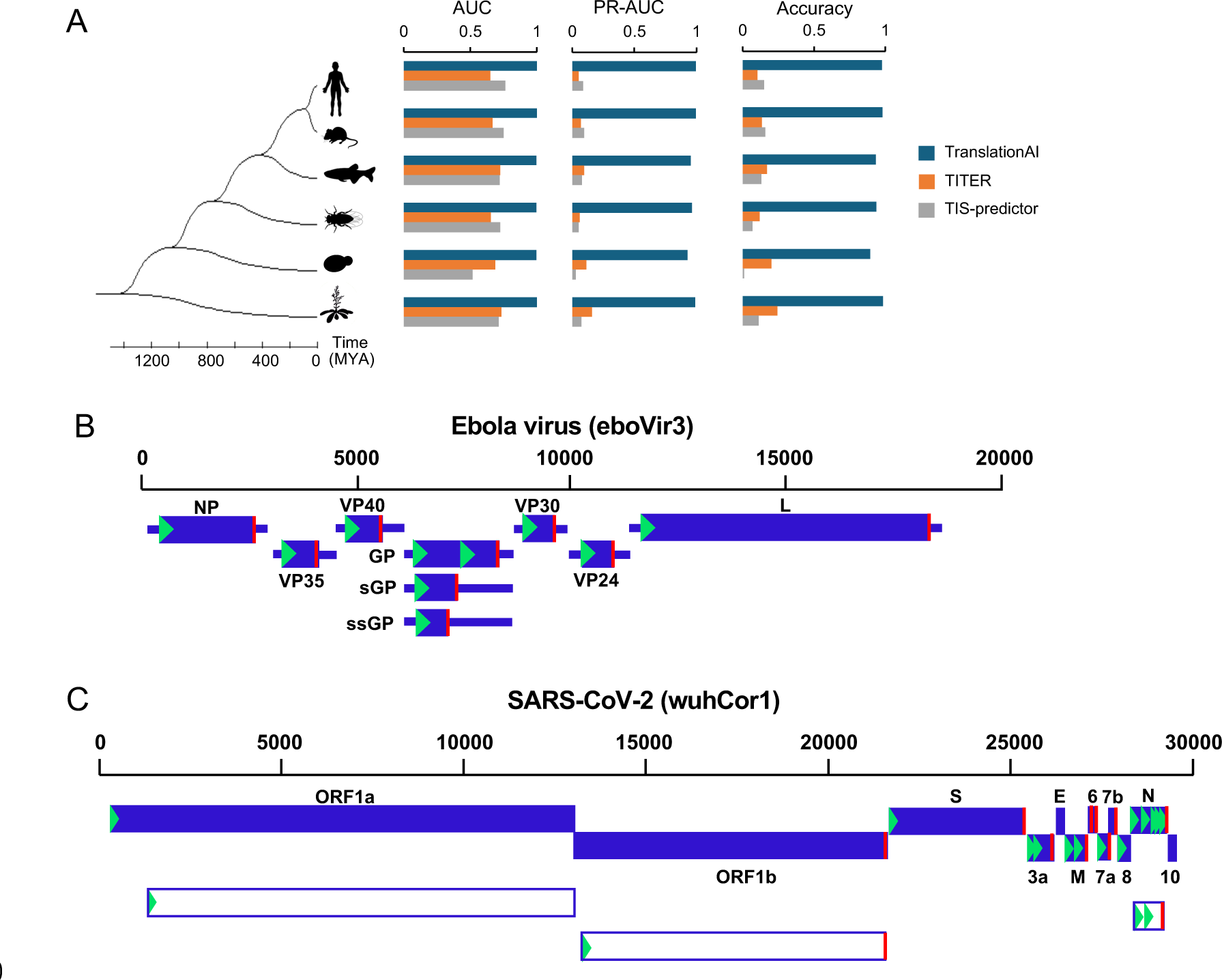
TranslationAI accurately predicts TIS/TTS of eukaryotes, prokaryotes, and viruses. **A.** The AUC, PR-AUC, and prediction accuracy of TIS prediction across tested eukaryotes (Human, Mouse, Zebrafish, Drosophila, Arabidopsis, and budding yeast *S. cerevisiae*). The predictions of two previous models are also included as a comparison. **B-C.** The prediction of TISs/TTSs on Ebola genomic or RNA sequences **B.** and SARS-CoV-2 genomic or subgenomic sequences **C.** The upper scales represent the genomic sequence position, the blue boxes indicate the annotated ORFs, and the white boxes indicate the newly predicted out-of-frame ORFs. The green triangles and red lines indicate the predicted in-frame TISs and TTSs, respectively.

In addition, we compared TranslationAI with two existing computational models, a deep learning model TITER(20) and a linear regression model TIS-predictor(23). Both previous models incorporated the surrounding local sequence as input to predict translation initiation sites, with TITER considering 200 nt and TIS-predictor considering 23 nt on both sides of TIS. Notably, we found that TranslationAI surpassed the other two models as judged by both the Area under the ROC Curve (AUC) and PR-AUC of TIS prediction across all tested eukaryotes (Fig. 4A and Table S1). The high performance of TranslationAI on canonical translation sites underscores the significant advance of our model over the previous ones, as the new model considers the entire mRNA rather than focusing solely on a limited region around candidate AUGs. Additionally, while TITER and TIS-predictor were primarily designed to predict non-canonical TIS by incorporating both AUG and near-cognate start codons, TranslationAI was specifically developed to predict translation sites without any prior bias toward non-canonical translation, and thus outperforms TITER and TIS-predictor in predicting canonical TISs that account for the majority of our training dataset.

We further assess the performance of TranslationAI model trained with human transcripts on the polycistronic transcription units of E. *coli*, which may reflect how much of the coding sequence might be conserved between eukaryotes and prokaryotes. Surprisingly, we obtained a 65% accuracy in *E. coli* with the deep learning model using all parameters trained on the human transcriptome (Fig. S5), even though most *E. coli* genes are transcribed as polycistronic units with very short UTRs and different set of genetic codons. Considering the lack of transcriptomic and proteomic information for most bacteria, this model could be useful in predicting their ORFs based solely on their genome. Furthermore, we extended this model on the transcriptomes of chloroplast (from *Arabidopsis*) and mitochondrion (from human), and the resulting prediction accuracies were ∼30% and <1%, respectively (Fig. S5). Given that the translation mechanism and the genetic codon in chloroplast and mitochondrion are drastically different from the human cells, such drop of prediction accuracy is not unreasonable.

Finally, we tested whether this model can predict the TISs/TTSs of human viruses that rely on the host cell machinery for protein synthesis, albeit through different translational mechanisms (35). Taking two RNA viruses (Ebola and SARS-CoV-2) as examples, we found that the model trained with human data can accurately predict all TISs and TTSs of Ebola virus even when we used the entire genomic RNA sequence as the input (Fig. 4B and Fig. S6A). In addition to the annotated TISs and TTSs, we also predicted a new internal translation start site in the GP gene in Ebola (Fig. 4B), suggesting an additional translation isoform for GP with truncated N-terminus.

Compared to Ebola virus, TranslationAI showed a lower prediction accuracy for SARS-CoV-2, with 8 out of 12 annotated TISs and 10 out of 12 annotated TTSs being predicted with high confidence by sub-genomic RNAs (36) (Fig. 4C and Fig. S6B). In addition, our model also made several false positive predictions of TISs/TTSs in SARS-CoV-2 genome (Fig. S6B). Similar performance was observed with either genomic or sub-genomic RNAs as input. A notable difference between these two viruses is that the Ebola genome is transcribed into monocistronic mRNAs with both UTRs, whereas the SARS-CoV-2 mRNAs are mainly polycistronic genomic or subgenomic RNAs with short spacer sequences between each ORF (36,37). This may partially explain the different performances of TranslationAI for different viruses.

Taken together, our results suggest that the model containing parameters trained on human transcriptome can accurately predict canonical ORFs in a wide range of organisms. However, in certain cases where the translation regulation is drastically different (e.g., short UTRs or different codon system), the application of this model should be carried out with caution.

## Discussion

As one of the most effective and powerful machine learning methods, the deep learning network has been widely used to study complex biological questions ranging from population genetics to precision medicine (12–15). Here we developed a deep learning architecture, TranslationAI, for *ab initio* prediction and systematic analysis of translation initiation and termination in entire transcriptome. Remarkably, the model trained with ∼70% of human transcripts achieved a near-perfect prediction accuracy in not only human, but also a wide range of eukaryotic organisms. The model can also identify new ORFs in annotated noncoding RNAs, some of which were confirmed by independent Ribo-seq and mass spectrometry experiments. Further analysis of this predictive model also identified a new regulation rule for translation termination, providing mechanistic insights into the complex regulation of RNA translation.

The final TranslationAI-2k model strategically utilized a flanking sequence of 1000 nt around the position of interest. We chose this design aiming to address the limitations observed in prior models that predominantly focused on characteristics of local sequences (20,23). By incorporating long-range sequence features, this model effectively captures potential interactions within mRNA sequences, including the long-range interactions of both UTRs, detecting frame shifts following one or two nucleotide deletions, and examination of codon bias within CDS.

The remarkable accuracy of TranslationAI suggests that this “black box” model have learned useful information with biological relevance. We found that the model has successfully learned the identity of canonical start and stop codons (Fig. 1F), which account for the vast majority (99.4% start codon and 99.6% stop codon) of the training set. The model did not identify any TIS/TTS with noncanonical start or stop (i.e, non-AUG start or non-UAG/UAA/UGA stop), probably because they are extremely rare in the training data. Additional features, including triplet reading frameshift, UTR sequences, and codon bias also contribute to the prediction accuracy. In addition, the long-range interactions contributed to the prediction accuracy of this model, as simultaneously disrupting the 5′-UTR and 3′-UTR cause synergistic effect in reducing the prediction scores (Fig. 1G). Moreover, certain internal sites in each mRNA also contributed to the TranslationAI prediction in a context-dependent fashion (Fig. S2B), however the underlying biological mechanisms are unclear. All these features contributed partially to the model in terms of TIS/TTS prediction, as disrupting these features did not completely invalidate the prediction (Fig. 1G-I).

Surprisingly, several known features associated with translation regulation, including the 3′-UTR sequence, synonymous mutation, and sequences of translation product, showed small effect on the accuracy of TIS/TTS prediction, implying that they may not play a dominant role in selection of translation sites. More importantly, we found several new features can influence the prediction of TIS/TTS, such as the G/C rich motif around the strong TTS, suggesting additional determinants for translation termination. These features may open up new avenues for investigating the molecular mechanisms underlying translation control. Future exploration of these predictive features may provide mechanistic insights into the functional interactions between the mRNA and translation machine.

We also used this model to assess the single nucleotide variants (SNVs) that may alter the TISs/TTSs and cause functional consequences. Surprisingly, we did not identify any disease-associated SNVs in the ORF that can alter the TIS/TTS prediction, unless the mutation directly changed the start or stop codon. This is consistent with the finding that the synonymous/non-synonymous substitutions of amino acids had little effect on TIS/TTS prediction (Fig. 1I). These results suggest that small changes in coding sequence do not significantly impact ORF selection by the translation machinery, *i.e.,* the translation initiation and termination appear to be unaffected by the elongation steps. Alternatively, it is also possible that the model does not rely on the nucleotide composition within ORFs for its prediction.

This model also allowed us to identify novel regulatory motifs associated with TIS and TTS. We discovered, for the first time, that the strong stop codons are surrounded by novel motifs that are generally CG-rich, especially in the third position of the codons immediately before the stop codon. Consistently, there is a codon bias associated with the strong termination site (Fig. 2E), and the synonymous mutations with low CG-contents will introduce translation readthrough in a reporter assay (Fig. 2F). Because the GC-rich codons were found to be associated with slow translation (38), we speculate that the strength of the stop codon is associated with the decreased translation elongation rate prior to reaching the stop codon. This regulation may be attributed to the changes in the strength of ribosome’s interaction with the mRNA before the ribosomes reach the termination site, which “decelerate” the translation before an efficient termination. Alternatively, the motif could alter the ribosome’s conformation, thereby influencing translation termination. This finding suggests that the translation regulation is more intricate than previously thought, and further exploration on this phenomenon may shed light on the mechanism of translation termination and readthrough.

The deep learning network trained with human transcripts achieved a surprisingly high accuracy for the prediction of ORFs in bacteria, yeast, plants and certain viruses using only their genome sequences as input (Fig. 4). This result is particularly interesting given that some of these organisms have polycistronic transcription units or genomes, suggesting that the TranslationAI can predict not only uORFs and dORFs in human, but also ORFs in polycistronic sequences from other organisms. However this model failed to predict mitochondrial ORFs and showed mediocre performance for SARS-CoV-2, probably due to the presence of frameshift regions, overlapping ORFs, and a lack of sufficient intergenic sequences to separate different ORFs. These results suggest that additional training may be required to improve the model’s performance on more complex or noncanonical genomes.

Despite its high accuracy in predicting canonical TISs, TranslationAI has some limitations. First, our model currently struggles in predicting non-canonical TISs, especially for uTISs with non-AUG start codons. This is probably due to the bias in the training dataset, which mainly consists of mono-cistronic translation units with AUG start codon. We have trained the model with non-AUG TIS, however distinguishing positive TISs from background proved challenging, probably due to the insufficient and noise experimental data for non-AUG start codon in human transcriptome. We also attempted to train this model using a subset of mRNAs with multiple annotated ORFs, however the accuracy (∼75%) was not as high as that of canonical transcripts, again possibly due to the small size of the training data.

Furthermore, although this model can predict the strengths of different TISs in the presence of multiple ORFs, it is still challenging to predict which TIS will be used in specific cell lines or tissues. Expanding the ribosome profiling data in different cell types may help refine the tissue-specific translation prediction model. Finally, the model uses a stringent threshold under the assumption that there is a dominant ORF in each transcript. While this assumption holds well for most transcripts, it may be incorrect in some cases. With more non-canonical ORFs being discovered, the score threshold may need to be modified to better fit different type of transcripts. In summary, with increasing translation datasets and more refined parameters, we believe that the TranslationAI framework will improve ORF prediction and deepen our understanding of the complexity in translation control.

## Methods

### Architecture of the TranslationAI deep learning model

We constructed a deep residual neural network using one-hot encoding to represent mRNA sequences as input (Fig. 1A). Our approach employed a 32-layer dilated convolutional neural network architecture (Fig. S1) to generate an output matrix representing the probabilities of a given position being a TIS, TTS, or neither (NS).

Four models were developed with distinct architectures, incorporating 60, 200, 600, or 2,000 nucleotides on both sides of a position of interest within the full-length mRNA sequence as input (Fig. S1). The segmentation of sequences into normalizing input sizes (60, 200, 600, or 2,000 nt) facilitates batch processing within the programming framework. Each model produces probability scores for TIS, TTS, and NS, with the sum of these probabilities equaling one.

The TranslationAI consists of several stacked residual blocks that connect the input layer to the penultimate layer, a framework previously developed for image recognition (39). A convolutional unit with softmax activation links the penultimate layer to the output layer. To enhance convergence speed during training, the output of every fourth residual block is added to the input of the penultimate layer (Fig. S1).

The fundamental unit of the TranslationAI model is a residual block, which contains two batch-normalization layers, two rectified linear units (ReLU), and two convolutional units arranged in a specific sequence (Fig. S1). Each residual block includes three hyper-parameters: N, W, and D. N represents the number of convolutional kernels, W signifies the window size, and D indicates the dilation rate of each convolutional kernel. Given that a convolutional kernel with window size W and dilation rate D processes features across (W-1)×D neighboring positions, a residual block with two convolutional units can handle features across 2(W-1)×D neighboring positions. As a result, a TranslationAI model with K stacked residual blocks is capable of extracting features spanning 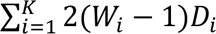 Neighboring positions, where *Ni*, *Wi*, and *Di* correspond to the hyper-parameters of the i-th residual block.

### Model training and testing

To obtain a comprehensive, well-curated, and non-redundant set of sequences, we utilized the RefSeq gene annotation and extracted 47,098 protein-coding transcripts with 47,098 TIS-TTS pairs. Different splicing isoforms of the same gene are treated as different transcripts. For model training, we selected transcripts located on chromosomes 2, 4, 6, 8, 10-22, X, and Y, which contain 13,707 genes with 34,292 TIS-TTS pairs. For model evaluation, the transcripts from chromosomes 1, 3, 5, 7, and 9, containing 5,524 genes with 12,806 TIS-TTS pairs, were used. To mitigate overfitting, 10% of the training set were randomly chosen for early-stopping determination during training, while the remainder were used for model training.

We employed a sequence-to-sequence approach with a chunk size of 6,000 for model training and testing. The full-length mature mRNA sequences were one-hot encoded, with A, C, G, and U mapped to [1, 0, 0, 0], [0, 1, 0, 0], [0, 0, 1, 0], and [0, 0, 0, 1], respectively. We then zero-padded the one-hot encoded nucleotide sequences until their lengths were multiples of 6,000 and further padded them with flanking sequences of length S/2 at the beginning and end, where S corresponds to 60, 200, 600, or 2,000 for the TranslationAI-60, TranslationAI-200, TranslationAI-600, and TranslationAI-2k models, respectively. The padded nucleotide sequences were divided into blocks of length S/2 + 6,000 + S/2, with the i-th block consisting of nucleotide positions from 6,000(i-1) – S/2 + 1 to 6,000i + S/2. In parallel, the output label sequences were one-hot encoded as TIS ([0, 1, 0]), TTS ([0, 0, 1]), or neither ([1, 0, 0]). The one-hot encoded TIS/TTS output label sequences were also zero-padded until their lengths reached multiples of 6,000, and then split into 6,000-length blocks, with the i-th block containing positions from 6,000(i-1) + 1 to 6,000i. These one-hot encoded nucleotide sequences and their corresponding label sequences served as the inputs and outputs for the models, respectively.

The models were trained for 10 epochs using a batch size of 18 on two NVIDIA Tesla K80 GPUs. During training, the categorical cross-entropy loss between target and predicted outputs was minimized using the Adam optimizer. The optimizer’s learning rate was set to 0.001 for the first six epochs and halved in each subsequent epoch. For each model, the training procedure was repeated 10 times, and the top five models based on performance were selected. During testing, each input was assessed using all five trained models, and the average of their outputs was utilized as the predicted output. This rigorous training and testing process aimed to ensure optimal performance and reliability in the resulting models.

The performance of the TIS/TTS prediction models was measured with top-k accuracy. This metric measures the fraction of correctly predicted positions among the top-k predicted sites, where k represents the number of true TIS/TTS sites in the dataset. For example, if the dataset contains k1 TISs and k2 TTSs, we set the score cutoff to predict exactly the top-k1 and top-k2 positions as TIS and TTS sites, respectively. It is worth noting that, unless specifically mentioned otherwise, only the top-k predicted sites were used for downstream analysis in our study.

During the development of the model, we have experimented with various hyper-parameters of this mode, including the input window size (60 nt, 200 nt, 600 nt, 2k and even 10k) and different convolutional filter sizes. However, it is worth noting that we did not systematically tune the model hyper-parameters. Therefore, the model hyper-parameters used here may not represent the optimal combinations. The results shown in the study probably represent the lower bound of this deep learning framework in the task of predicting TIS and TTS.

### Role of transcript lengths in mRNA translation

To study the effect of transcript length on translation sites prediction, we sorted the mRNAs in order of increasing length. Using the TranslationAI-200 and TranslationAI-2k models, we calculated the positive rate of TIS/TTS for each transcript by comparing the number of positive TISs to the total number of transcripts with lengths shorter than the given transcript. We plotted these positive rates against the transcript length, as shown in Fig. 1C. Additionally, we included a histogram of transcript lengths from the 47,098 transcripts analyzed in the background of the figure.

### Perturbations of synonymous mutation

To investigate the impact of codon bias on predictions, we introduced perturbations by substituting codons with synonymous counterparts. Three scenarios were considered: 1) Random Synonymous Substitutions: Codons were randomly replaced with synonymous counterparts. 2) Higher CAI Substitutions: Codons were replaced with synonymous counterparts possessing a higher CAI. The Relative Synonymous Codon Usage (RSCU) for each codon was obtained from the Codon Statistics Database (http://codonstatsdb.unr.edu). The RSCU values were calculated based on genes encoding ribosomal proteins, known for their high expression and consequent strong codon bias (40). 3) Lower CAI Substitutions: Codons were replaced with synonymous counterparts possessing a lower CAI, following the same procedure as described above.

### Occlusion sensitivity analysis

To assess the feature importance of individual positions in predicting TIS and TTS, we performed the occlusion sensitivity analysis, which provides a clear visualization of the contribution of each nucleotide to the scoring of TIS and TTS in an mRNA sequence. We masked each nucleotide in the mRNA with “N” one by one, and calculated the resulting score change of the predicted TIS/TTS.

### Codon distribution at upstream of stop codon

To examine the codon distribution upstream of stop codon, we quantified the frequency of each codon at the -30nt position relative to both strong and weak stop codons. Specifically, we calculated the frequency of each codon as the number of occurrences of that codon at the -30nt position divided by the total number of RNAs analyzed at that position.

### Conservation analysis

To evaluate the conservation levels of different RNAs, we obtained the PhastCons46 scores from the PhastCons46 or PhyloP46 track in the UCSC Genome Browser. These scores were generated from genome alignments of 46 vertebrates, including human (hg19), and provide a measure of evolutionary conservation across species. Each box plot displays the median as the central line, while the upper and lower edges of the box represent the first and third quartiles, respectively.

### Non-canonical TISs/TTSs identification

To systematically identify non-canonical TISs and TTSs in transcriptomes, we input human transcriptome into TranslationAI model and retained only those RNAs that met the threshold criteria (prediction score > 0.5) for both TIS and TTS as translatable RNAs. The ncRNA dataset were retrieved from Ensembl build 75, which was previously used in the published study (8) that identified hundreds of noncanonical ORFs using ribosome profiling and mass-spectrometry.

To compare the TISs predicted by TranslationAI with TISs identified by ribosome profiling, we expanded our predicted TIS library by retaining the top 10 TISs for each gene predicted by the TranslationAI. This approach was necessary as ribosome profiling often identifies multiple ORFs from one gene.

The alternative TISs were defined as the TIS with the highest score (>0.1), excluding the annotated TIS. In addition, in-frame alternative TISs were defined as TISs where the distance between the annotated TIS and alternative TIS is a multiple of three.

### Ribosome footprint data analysis

The ribosome profiling data analysis was performed on a previously published ribosome footprint data that were retrieved from no drug treatment iPSC and iPSC-induced iPSC-derived cardiomyocytes cells being deposited in GEO database under accession number GSE131650. The genome assembly used throughout this manuscript is hg19/GRCh37 annotated by GENCODE v27. We first filtered out reads aligning to rRNAs using Bowtie v1.0.0 and aligned the remaining reads to the annotated transcriptome with STAR v2.7.5a, using the --outSJfilterReads Unique -- outFilterMultimapNmax 1 --outFilterMismatchNmax 999 -- outFilterMismatchNoverReadLmax 0.04 to filter multiple aligned and redundant reads. These alignments were assigned a specific P-site nucleotide using a 12-nt offset from the 3′ end of reads. Metagene plots were generated by normalizing read density around the start and stop codons of each gene (i.e, read depth at each position divided by the average reads of each position in one transcript and the number of the covered transcripts).

To estimate the correlation between predicted TIS/TTS scores and mRNA translation efficiency, we ranked and divided the scores into percentiles (n∼1580 transcripts/percentile, Fig. S2G-S2H). We calculated the translation efficiency of transcripts from ribosome profiling data in two human cell lines as the ratio of Ribo-seq read density to RNA-seq read density.

### Motif identification of strong and weak TISs/TTSs

Motif discovery and analysis were performed using the Meme suite (v5.0.5). The strong and weak TIS/TTS containing sequences (-30nt : + 33nt) were collected as primary sequences for motif prediction. Additionally, all TIS/TTS containing sequences (-30nt : +33nt) were used as control sequences. Meme was employed to identify enriched motifs, with a maximum length of 63nt, that occur any number of times within the given sequence.

### MS Proteomics and HLA Peptidomics data analysis

MS-based proteomics data are deposited to the ProteomeXchange Consortium via the Proteomics Identifications Database (PRIDE) partner repository with the dataset identifier PXD014031 and PXD000394. Using an open search engine pFind (v3.1.3), we searched the two previously published human proteome datasets against a combined database (25692 proteins) containing all UniProt human proteins and the potential non-canonical ORFs-coded peptides (uORFs, dORFs, dual frame ORFs, and new ORFs). We selected positive mass spectra using following thresholds: q < 0.01, peptides length ≥ 8, missed cleavage sites ≤ 3, allowing only common modifications (cysteine carbamidomethylation, oxidation of methionine, protein N-terminal acetylation, pyro-glutamate formation from glutamine, and phosphorylation of serine, threonine, and tyrosine residues).

The resulting peptides were searched against non-redundant human protein database using blastp-short and the peptides with less than two mismatches from known proteins were removed.

### Transcriptomes of other species

The genome sequence and gene annotations of *Mus musculus* (mm10), *Danio rerio* (danRer11), *Drosophila melanogaster* (dm6), Ebola virus (eboVir3), SARS-CoV-2 (wuhCor1), and human mitochondrion are obtained from UCSC database. The sub-genomic sequences of SARS-CoV2 were obtained from a published study (36). The genome sequence and gene annotation of Arabidopsis thaliana (TAIR10) are obtained from TAIR database. Regarding E. coli (Escherichia coli K-12 substr. MG1655), its transcription unit is retrieved from EcoCyc. The mRNA information (5′-UTR, CDS, and 3′-UTR) for yeast was obtained from *Saccharomyces* Genome Database (SGD). The genome sequence and gene annotation of Chloroplast (Arabidopsis thaliana) was obtained from NCBI database. The genome sequence of human mitochondria was obtained from NCBI database and its gene annotation file was obtained from GENCODE database.

### Model comparison

TITER(20) (Translation Initiation siTE detectoR) is a deep learning-based framework designed to predict noncanonical TISs by incorporating both AUG and near-cognate start codons using high-throughput sequencing data. With the input of each TIS with its 200 nt flanking nucleotides, TITER outputs the probability of translation initiation for the given sequence. The source code for TITER was downloaded from https://github.com/zhangsaithu/titer. The TIS-predictor(23) is a machine learning model that was trained on instances of AUG and near-cognate start codon. The algorithm examines each codon and its 20 flanking nucleotides (23 nucleotides in total) to simulate and compute its probability score as a translation initiation site. The source code for TIS-predictor was downloaded from https://github.com/Agleason1/TIS-Predictor.

To evaluate the performance of TITER, TIS-predictor, and TranslationAI on canonical TIS AUG, we calculated the AUC, PR-AUC, and accuracy. For each RNA sequence input, we computed the probability score for all AUG codons, predicting the AUG with the highest score as the translation initiation site for each mRNA. These tools were tested on datasets from multiple species, including a Human test dataset and the whole transcriptomes of Mouse, Zebrafish, Drosophila, Arabidopsis, and the budding yeast (*S. cerevisiae*).

### Plasmid construction and Transfection

To construct the reporter (pcDNA5-HA-GFP-stop_motif-Nluc-Flag) for testing the translation readthrough products, the Fluc-Nluc luciferase sequence (a gift from prof. Rachel Green) was cloned into pcDNA5 backbone digested by SpeI and XbaI (41). The GFP sequence was amplified (fused with two BsmBI restriction fragments) and inserted before Nluc. The modified VDR sequences were synthesized (GENEWIZ) and cloned into BsmBI digested vector.

HEK293T cell line was cultured in DMEM (high glucose) medium containing 10% fetal bovine serum (FBS, Hyclone). To transient transfect plasmids into cells, 2 μg of reporters were transfected into cells in 6 well plate using lipofectamine 3000 (Invitrogen) according to the manufacturer’s instruction. After 48 h, cells were collected for further analysis of protein level.

### Western blot

Cells were lysed in laemmli buffer, and the total cell lysates were resolved with 4-20% ExpressPlus™ PAGE Gel (GeneScript). The following antibodies were used: HA-Tag antibody (CST: 3724S) was diluted by 1:2000, V5 antibody (CST: 13202S) was diluted by 1:2000, Flag antibody (Sigma: F1804-1MG) was diluted by 1:2000. The HRP-linked secondary antibodies (CST: 7076S) were used by 1:4000 dilution and the blots were visualized with the ECL reagents (Bio-Rad).

## Data and software availability

Training and testing data, prediction scores for all possible single nucleotide substitutions in the reference genome and source code are publicly hosted at GitHub (https://github.com/rnasys/TranslationAI). Prediction scores and source code are publicly released under GPL v3 and are free for use for academic and non-commercial applications.

## Acknowledgments

We thank Dr. Rachel Green for kindly sharing the plasmids of translation read-through and offering insightful suggestions on the potential mechanisms of translation termination. This work is supported by the Strategic Priority Research Program of Chinese Academy of Sciences (XDB38040100 to Z.W.), the National Key Research and Development Program of China (2021YFA1300503 and 2018YFA0107602 to Z.W.), the National Natural Science Foundation of China ( 32030064, 32250013, 91940303 and 31730110 to Z.W.; 32100430 to X.F.), the Starry Night Science Fund at Shanghai Institute for Advanced Study of Zhejiang University (SN-ZJU-SIAS-009) and the NIH Basic Research in Cancer Health Disparities Award to S.R.P. and J.A.F. (R01CA220314). Z.W. is also supported by CAS Interdisciplinary Innovation Team and the type A CAS Pioneer 100-Talent program.

## Author Contributions

Conceptualization, Z.W., X.F. and T.C.; Methodology, X.F., T.C., and Z.W.; Software, X.F. and T.C.; Experiments, C.C.; Writing and discussion, X.F., Z.W., T.C., and M.H.; Funding acquisition, X.F., and Z.W.

## References

1. Frankish, A., Diekhans, M., Ferreira, A.M., Johnson, R., Jungreis, I., Loveland, J., Mudge, J.M., Sisu, C., Wright, J., Armstrong, J. et al. (2019) GENCODE reference annotation for the human and mouse genomes. Nucleic Acids Res, 47, D766–D773.

2. Mouilleron, H., Delcourt, V. and Roucou, X. (2016) Death of a dogma: eukaryotic mRNAs can code for more than one protein. Nucleic Acids Res, 44, 14–23.

3. Hinnebusch, A.G. (2011) Molecular mechanism of scanning and start codon selection in eukaryotes. Microbiol Mol Biol Rev, 75, 434–467, first page of table of contents.

4. McNair, K., Ecale Zhou, C.L., Souza, B., Malfatti, S. and Edwards, R.A. (2021) Utilizing Amino Acid Composition and Entropy of Potential Open Reading Frames to Identify Protein-Coding Genes. Microorganisms, 9.

5. Jackson, R., Kroehling, L., Khitun, A., Bailis, W., Jarret, A., York, A.G., Khan, O.M., Brewer, J.R., Skadow, M.H., Duizer, C. et al. (2018) The translation of non-canonical open reading frames controls mucosal immunity. Nature, 564, 434–438.

6. Matsumoto, A., Pasut, A., Matsumoto, M., Yamashita, R., Fung, J., Monteleone, E., Saghatelian, A., Nakayama, K.I., Clohessy, J.G. and Pandolfi, P.P. (2017) mTORC1 and muscle regeneration are regulated by the LINC00961-encoded SPAR polypeptide. Nature, 541, 228–232.

7. Yang, Y., Fan, X., Mao, M., Song, X., Wu, P., Zhang, Y., Jin, Y., Yang, Y., Chen, L.L., Wang, Y. et al. (2017) Extensive translation of circular RNAs driven by N(6)-methyladenosine. Cell Res, 27, 626–641.

8. Chen, J., Brunner, A.D., Cogan, J.Z., Nunez, J.K., Fields, A.P., Adamson, B., Itzhak, D.N., Li, J.Y., Mann, M., Leonetti, M.D. et al. (2020) Pervasive functional translation of noncanonical human open reading frames. Science, 367, 1140–1146.

9. Raj, A., Wang, S.H., Shim, H., Harpak, A., Li, Y.I., Engelmann, B., Stephens, M., Gilad, Y. and Pritchard, J.K. (2016) Thousands of novel translated open reading frames in humans inferred by ribosome footprint profiling. Elife, 5.

10. Jaganathan, K., Kyriazopoulou Panagiotopoulou, S., McRae, J.F., Darbandi, S.F., Knowles, D., Li, Y.I., Kosmicki, J.A., Arbelaez, J., Cui, W., Schwartz, G.B. et al. (2019) Predicting Splicing from Primary Sequence with Deep Learning. Cell, 176, 535–548 e524.

11. Jumper, J., Evans, R., Pritzel, A., Green, T., Figurnov, M., Ronneberger, O., Tunyasuvunakool, K., Bates, R., Zidek, A., Potapenko, A. et al. (2021) Highly accurate protein structure prediction with AlphaFold. Nature, 596, 583–589.

12. Eraslan, G., Avsec, Z., Gagneur, J. and Theis, F.J. (2019) Deep learning: new computational modelling techniques for genomics. Nat Rev Genet, 20, 389–403.

13. Bogard, N., Linder, J., Rosenberg, A.B. and Seelig, G. (2019) A Deep Neural Network for Predicting and Engineering Alternative Polyadenylation. Cell, 178, 91–106 e123.

14. Zhou, J., Theesfeld, C.L., Yao, K., Chen, K.M., Wong, A.K. and Troyanskaya, O.G. (2018) Deep learning sequence-based ab initio prediction of variant effects on expression and disease risk. Nat Genet, 50, 1171–1179.

15. Vamathevan, J., Clark, D., Czodrowski, P., Dunham, I., Ferran, E., Lee, G., Li, B., Madabhushi, A., Shah, P., Spitzer, M. et al. (2019) Applications of machine learning in drug discovery and development. Nat Rev Drug Discov, 18, 463–477.

16. Sonenberg, N. and Hinnebusch, A.G. (2009) Regulation of translation initiation in eukaryotes: mechanisms and biological targets. Cell, 136, 731–745.

17. Lee, S., Liu, B., Lee, S., Huang, S.X., Shen, B. and Qian, S.B. (2012) Global mapping of translation initiation sites in mammalian cells at single-nucleotide resolution. Proc Natl Acad Sci U S A, 109, E2424–2432.

18. Gao, X., Wan, J., Liu, B., Ma, M., Shen, B. and Qian, S.B. (2015) Quantitative profiling of initiating ribosomes in vivo. Nat Methods, 12, 147–153.

19. Chen, W., Feng, P.M., Deng, E.Z., Lin, H. and Chou, K.C. (2014) iTIS-PseTNC: a sequence-based predictor for identifying translation initiation site in human genes using pseudo trinucleotide composition. Anal Biochem, 462, 76–83.

20. Zhang, S., Hu, H., Jiang, T., Zhang, L. and Zeng, J. (2017) TITER: predicting translation initiation sites by deep learning. Bioinformatics, 33, i234–i242.

21. Goel, N., Singh, S. and Aseri, T.C. (2020) Global sequence features based translation initiation site prediction in human genomic sequences. Heliyon, 6, e04825.

22. Kalkatawi, M., Magana-Mora, A., Jankovic, B. and Bajic, V.B. (2019) DeepGSR: an optimized deep-learning structure for the recognition of genomic signals and regions. Bioinformatics, 35, 1125–1132.

23. Gleason, A.C., Ghadge, G., Chen, J., Sonobe, Y. and Roos, R.P. (2022) Machine learning predicts translation initiation sites in neurologic diseases with nucleotide repeat expansions. PLoS One, 17, e0256411.

24. Leppek, K., Das, R. and Barna, M. (2018) Functional 5’ UTR mRNA structures in eukaryotic translation regulation and how to find them. Nat Rev Mol Cell Biol, 19, 158–174.

25. Mayr, C. (2019) What Are 3’ UTRs Doing? Cold Spring Harb Perspect Biol, 11.

26. Fields, A.P., Rodriguez, E.H., Jovanovic, M., Stern-Ginossar, N., Haas, B.J., Mertins, P., Raychowdhury, R., Hacohen, N., Carr, S.A., Ingolia, N.T. et al. (2015) A Regression-Based Analysis of Ribosome-Profiling Data Reveals a Conserved Complexity to Mammalian Translation. Mol Cell, 60, 816–827.

27. Kozak, M. (1987) An analysis of 5’-noncoding sequences from 699 vertebrate messenger RNAs. Nucleic Acids Res, 15, 8125–8148.

28. Kozak, M. (1986) Point mutations define a sequence flanking the AUG initiator codon that modulates translation by eukaryotic ribosomes. Cell, 44, 283–292.

29. Loughran, G., Jungreis, I., Tzani, I., Power, M., Dmitriev, R.I., Ivanov, I.P., Kellis, M. and Atkins, J.F. (2018) Stop codon readthrough generates a C-terminally extended variant of the human vitamin D receptor with reduced calcitriol response. J Biol Chem, 293, 4434–4444.

30. Mills, E.W., Wangen, J., Green, R. and Ingolia, N.T. (2016) Dynamic Regulation of a Ribosome Rescue Pathway in Erythroid Cells and Platelets. Cell Rep, 17, 1–10.

31. de Klerk, E. and t Hoen, P.A. (2015) Alternative mRNA transcription, processing, and translation: insights from RNA sequencing. Trends Genet, 31, 128–139.

32. Manske, F., Ogoniak, L., Jurgens, L., Grundmann, N., Makalowski, W. and Wethmar, K. (2023) The new uORFdb: integrating literature, sequence, and variation data in a central hub for uORF research. Nucleic Acids Res, 51, D328–D336.

33. Wan, J. and Qian, S.B. (2014) TISdb: a database for alternative translation initiation in mammalian cells. Nucleic Acids Res, 42, D845–850.

34. Huang, J.Z., Chen, M., Chen, D., Gao, X.C., Zhu, S., Huang, H., Hu, M., Zhu, H. and Yan, G.R. (2017) A Peptide Encoded by a Putative lncRNA HOXB-AS3 Suppresses Colon Cancer Growth. Mol Cell, 68, 171–184 e176.

35. Stern-Ginossar, N., Thompson, S.R., Mathews, M.B. and Mohr, I. (2019) Translational Control in Virus-Infected Cells. Cold Spring Harb Perspect Biol, 11.

36. Wang, D., Jiang, A., Feng, J., Li, G., Guo, D., Sajid, M., Wu, K., Zhang, Q., Ponty, Y., Will, S. et al. (2021) The SARS-CoV-2 subgenome landscape and its novel regulatory features. Mol Cell, 81, 2135–2147 e2135.

37. Banerjee, A.K., Blanco, M.R., Bruce, E.A., Honson, D.D., Chen, L.M., Chow, A., Bhat, P., Ollikainen, N., Quinodoz, S.A., Loney, C. et al. (2020) SARS-CoV-2 Disrupts Splicing, Translation, and Protein Trafficking to Suppress Host Defenses. Cell, 183, 1325–1339 e1321.

38. Tunney, R., McGlincy, N.J., Graham, M.E., Naddaf, N., Pachter, L. and Lareau, L.F. (2018) Accurate design of translational output by a neural network model of ribosome distribution. Nat Struct Mol Biol, 25, 577–582.

39. He, K., Zhang, X., Ren, S. and Sun, J. (2016) Deep Residual Learning for Image Recognition. Proceedings of the IEEE Conference on Computer Vision and Pattern Recognition, 770–778.

40. Subramanian, K., Payne, B., Feyertag, F. and Alvarez-Ponce, D. (2022) The Codon Statistics Database: A Database of Codon Usage Bias. Mol Biol Evol, 39.

41. Wangen, J.R. and Green, R. (2020) Stop codon context influences genome-wide stimulation of termination codon readthrough by aminoglycosides. Elife, 9.

