## Supplementary Figures for "Analysis of RNA translation with a deep learning architecture provides new insight into translation control"

**Summary of supplemental information**

Supplementary Figure 1. Architectures of TranslationAI-60, TranslationAI-200, TranslationAI-600, and TranslationAI-2k.

Supplementary Figure 2. Key features learned during AI training for accurate prediction.

Supplementary Figure 3. Sequence features of strong and weak TISs/TTSs.

Supplementary Figure 4. Prediction of translatable non-coding RNA.

Supplementary Figure 5. Overlaps of predicted TIS/TTS and annotated TIS/TTS in Arabidopsis Chloroplast and human mitochondrion.

Supplementary Figure 6. All predicted TISs and TTSs of Ebola and SARS-CoV-2.

Supplementary Table 1. Comparisons of models' features and performances

Supplementary Table 2. Predictions of *in silico* perturbations on different regions of mRNAs

Supplementary Table 3. Primers and synthesized sequences

Supplementary Table 4. The newly predicted uORFs, dORFs, and overlapping ORFs with different reading frames.

Supplementary Table 5. Predicted translatable lncRNA

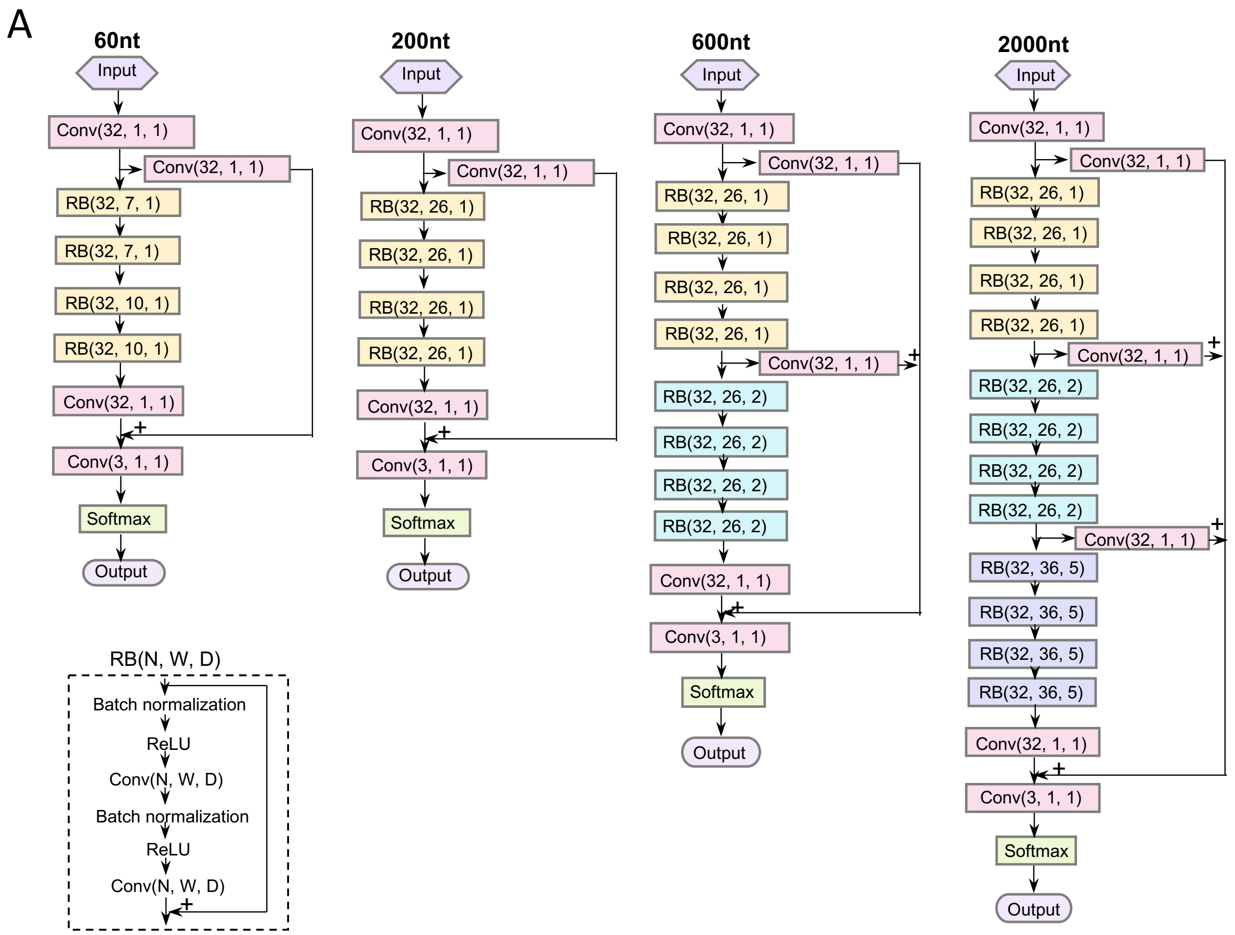


**Figure S1. Architectures of TranslationAI-60, TranslationAI-200, TranslationAI-600, and TranslationAI-2k.** Each architecture takes as input flanking nucleotide sequences of different lengths (60, 200, 600, and 2,000 nucleotides) on each side of a given position and outputs the probability of the position being a translation initiation site (TIS), translation termination site (TTS), or neither. The architectures are primarily composed of convolutional layers Conv(N, W, D), where N, W, and D denote the number of convolutional kernels, window size, and dilation rate of each kernel in the layer, respectively.

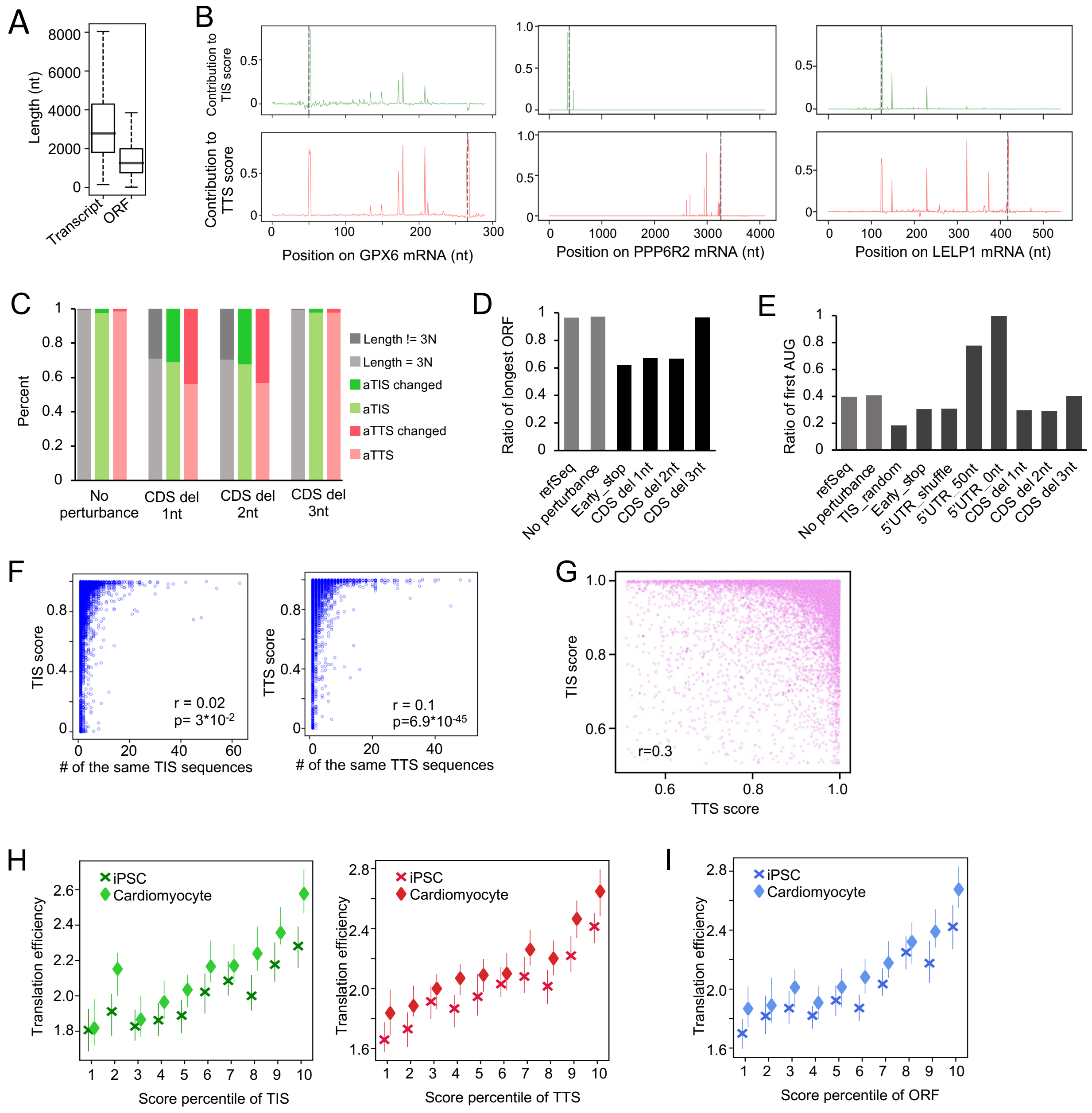


**Figure S2. Key features learned during AI training to achieve accurate prediction of TISs and TTSs. A.** Boxplot analysis of transcript length and ORF length from refGene. **B.** Visualization of the contribution of each nucleotide to the scoring of TIS and TTS (occlusion sensitivity analysis) in randomly selected mRNAs GPX6, PPP6R2, and LELP1. The TISs and TTSs of each mRNA were marked by grey dotted line. **C.** Percentage of the transcripts using annotated TIS/TTS, changed TIS/TTS, and 3N rule after introducing of frameshift permutations by deleting 1nt, 2nt, and 3nt (no frameshift). **D.** Ratio of longest ORF after introducing premature stop codons or frameshifts (as in panel A). **E.** Ratio of first AUG usage affected by many permutations: changes in TIS, introducing early stop codon, shuffling of 5′-UTR sequences, shortening of 5′-UTR, and introducing frameshift. **F.** Correlation analysis (Spearman's correlation coefficient) between the predicted score of TIS/TTS and the number of transcript isoforms containing the same TIS/TTS sequences. **G.** Correlation analysis (Spearman's correlation coefficient) between the TIS and TTS scores in the same ORF. **H.** The TIS/TTS scores predicted by TranslationAI are positive correlated with the translation efficiency in both iPSC and iPSC-derived cardiomyocytes cells. The total TIS/TTS scores were ranked and equally divided into 10 bins (10 percentiles, each containing ~1580 transcripts). The translation efficiency estimated by the ratio of read density from Ribosome profiling (Ribo-seq) normalized by mRNA abundance determined by RNA-seq. **I.** The ORF scores exhibit a positive correlation with the translation efficiency of corresponding transcripts. Each transcript's ORF score was determined by averaging the TIS and TTS scores obtained from TranslationAI. Similar to panel G of Figure S2, the score percentiles of ORF were calculated and divided into 10 bins.

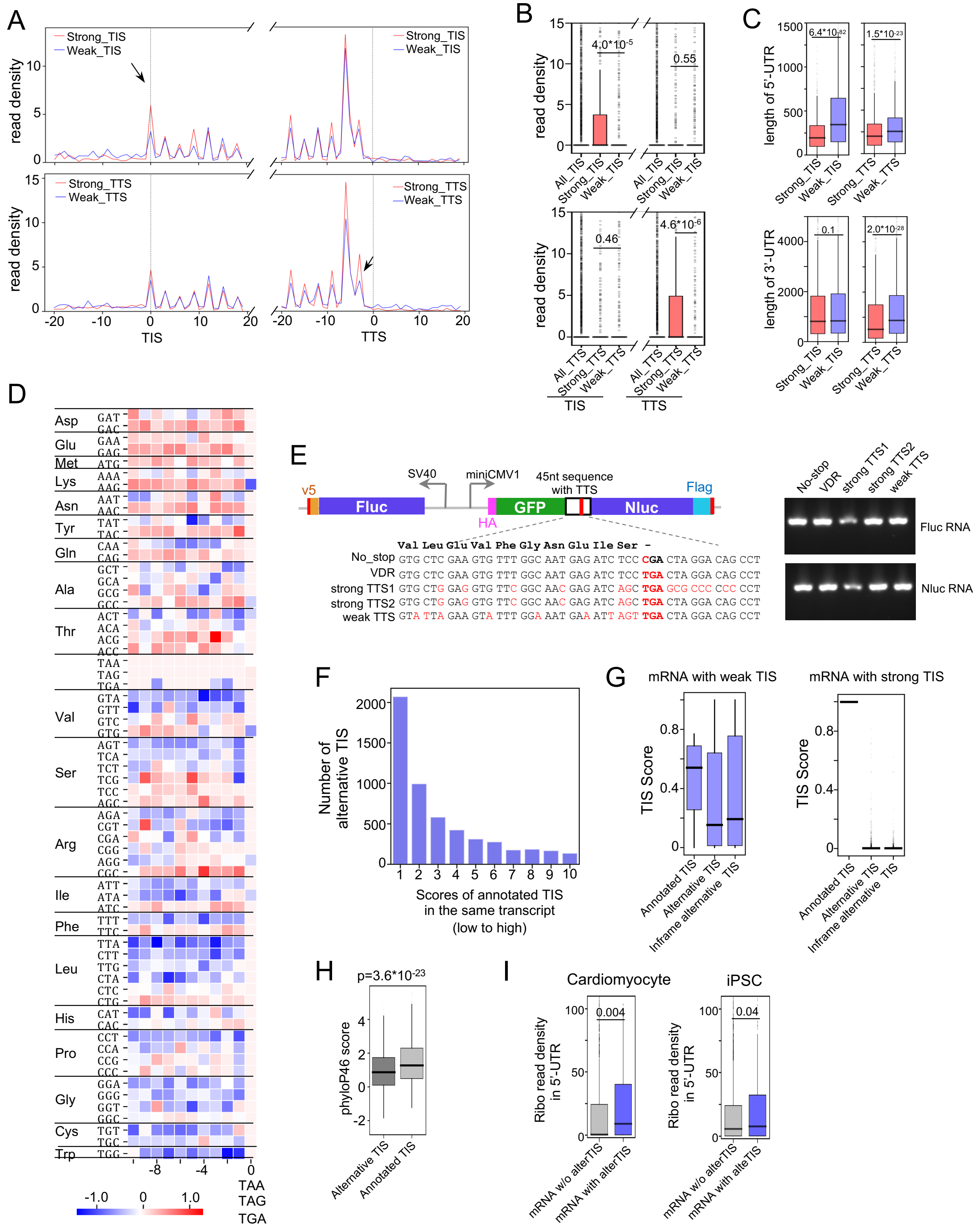

A
Strong_TIS
Weak_TIS
read density
TIS
Strong_TTS
Weak_TTS
TTS
B
read density
4.0*10-5
0.55
0.46
4.6*10-6
All_TIS
Strong_TIS
Weak_TIS
All_TTS
Strong_TTS
Weak_TTS
TIS
TTS
C
length of 5'-UTR
length of 3'-UTR
6.4*10-82
1.5*10-23
0.1
2.0*10-28
D
Asp
Glu
Met
Lys
Asn
Tyr
Gln
Ala
Thr
Val
Ser
Arg
Ile
Phe
Leu
His
Pro
Gly
Cys
Trp
TAA
TAG
TGA
E
v5
Fluc
SV40
miniCMV1
HA
GFP
45nt sequence with TTS
Nluc
Flag
Val Leu Glu Val Phe Gly Asn Glu Ile Ser -
No_stop GTG CTC GAA GTG TTT GGC AAT GAG ATC TCC CGA CTA GGA CAG CCT
VDR GTG CTC GAA GTG TTT GGC AAT GAG ATC TCC TGA CTA GGA CAG CCT
strong TTS1 GTG CTG GAG GTG TTC GGC AAC GAG ATC AGC TGA GCG CCC CCC CCT
strong TTS2 GTG CTG GAG GTG TTC GGC AAC GAG ATC AGC TGA CTA GGA CAG CCT
weak TTS GTA TTA GAA GTA TTT GGA AAT GAA ATT AGT TGA CTA GGA CAG CCT
No-stop
VDR
strong TTS1
strong TTS2
weak TTS
Fluc RNA
Nluc RNA
F
Number of alternative TIS
Scores of annotated TIS in the same transcript (low to high)
G
mRNA with weak TIS
mRNA with strong TIS
TIS Score
Annotated TIS
Alternative TIS
Inframe alternative TIS
H
p=3.6*10-23
phyloP46 score
Alternative TIS
Annotated TIS
I
Cardiomyocyte
iPSC
Ribo read density in 5'-UTR
0.004
0.04
mRNA w/o alterTIS
mRNA with alterTIS

**Figure S3. Sequence features of strong and weak TISs/TTSs. A.** Metagene analysis and **B.** boxplot depict the reads density on transcripts containing strong (red) and weak (blue) TIS/TTS (-20nt : +20nt) in iPSC-induced Cardiomyocyte cell line. **C.** Length distribution of 5′-UTR and 3′-UTR among the transcripts with strong and weak TIS/TTS. **D.** Heatmap of codon frequency at the -30nt position relative to both strong and weak stop codons, where the color bar shows the log10 fold change of each codon at the -30nt position between strong and weak stop codons. **E.** The corresponding RNA levels of Fluc and Nluc are shown for Fig. 2F. Different variants are described: No_stop: stop codon of VDR (TGA) mutated to CGA; strong TTS1: stop codon of VDR mutated to a strong TTS by changing the upstream 27nt and downstream 12nt sequences; strong TTS2: stop codon of VDR mutated to a strong TTS by changing the upstream 27nt sequence; weak TTS: stop codon of VDR mutated to a weak TTS by changing the upstream 27nt. See supplementary information in Table S2. **F.** Number of predicted alternative TISs of transcripts in each bin of 10% transcripts equally divided according to scores of annotated TIS, from low to high. **G.** Boxplots display predicted scores of annotated TIS, alternative TIS, and in-frame alternative TIS from transcripts containing strong and weak TIS/TTS. **H.** Conservation analysis of alternative TISs and annotated TISs in transcripts containing weak TISs. **I.** Boxplot analysis of reads density (Ribo-seq) on 5′-UTRs from transcripts with strong/weak TIS/TTS in iPSC cell line and iPSC-induced Cardiomyocyte cell line, respectively.

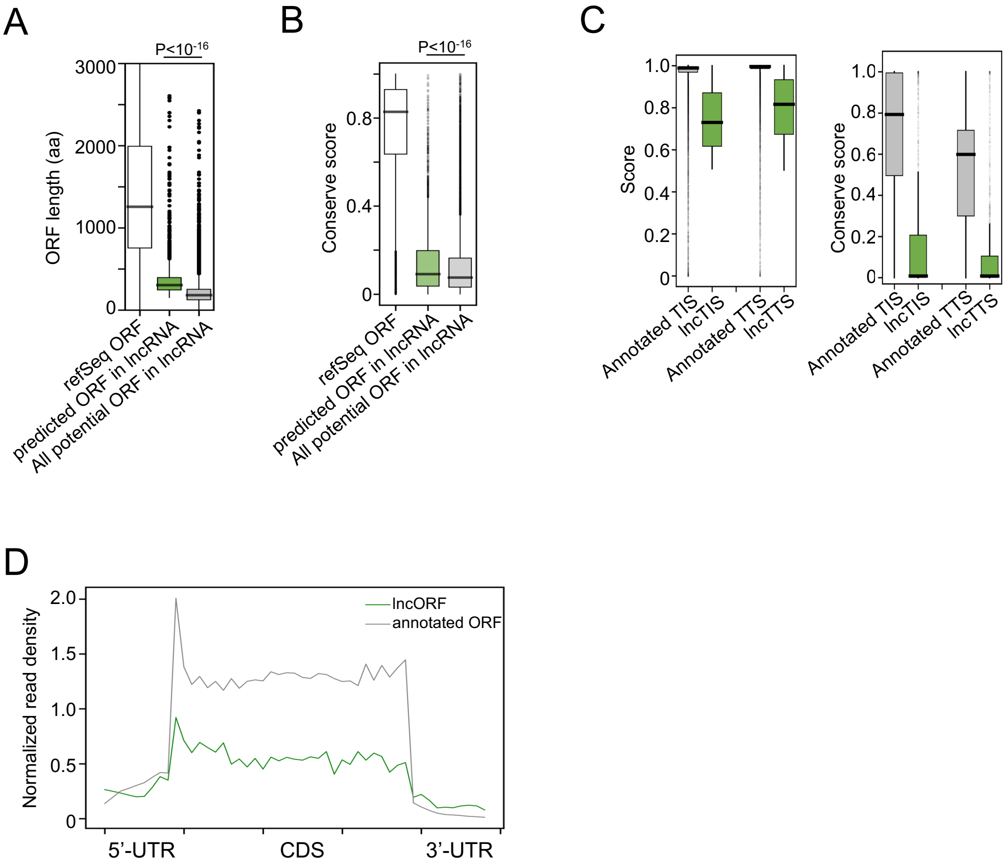


**Figure S4. Features of translatable non-coding RNA.** The distribution of **A.** ORF length and **B.** conservation are depicted for refSeq ORFs, predicted ORFs in lncRNA by translationAI, and all potential ORFs in lncRNA (defined as the longest fragments between ATG and TAA/TAG/TGA in all lncRNAs). P-values were calculated using Mann–Whitney U test. **C.** Prediction score and conservation analysis are shown for predicted TIS/TTS in lncRNA (lncTIS/lncTTS) and mRNA (Annotated TIS/Annotated TTS). **D.** Metagene analysis of Ribo-seq read density among translatable ncRNAs and control RNAs (mRNAs with a similar distribution of ORF lengths as predicted ORFs from lncRNAs).

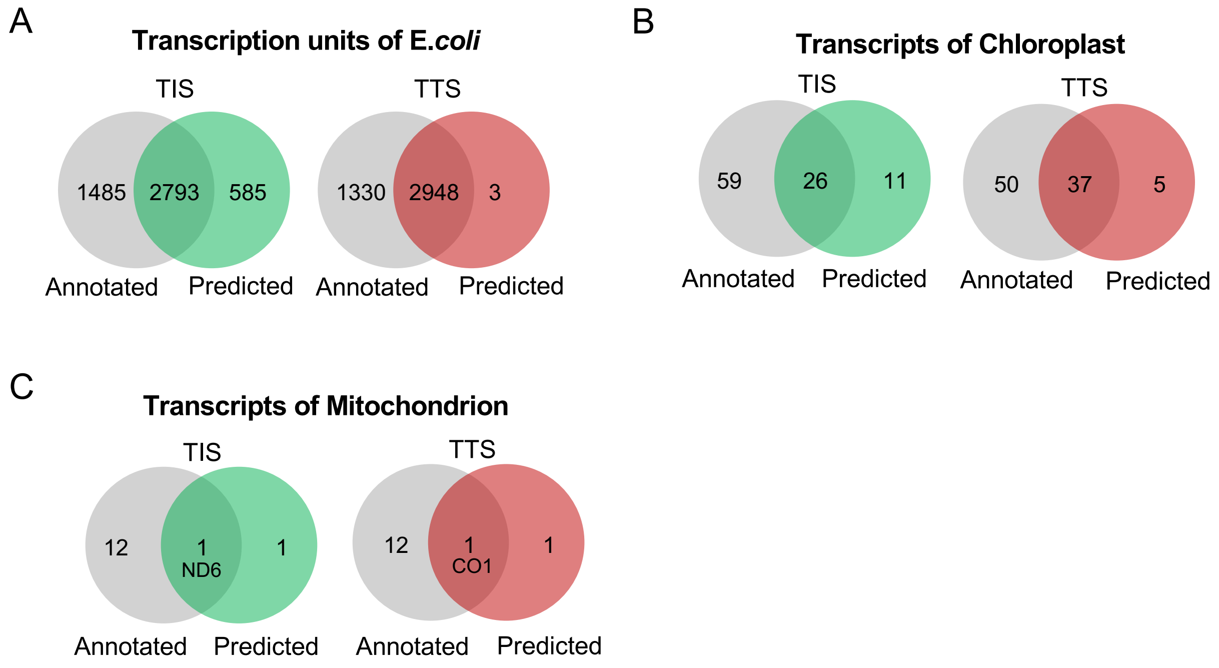


**Figure S5. Overlaps between predicted TIS/TTS and annotated TIS/TTS in Arabidopsis Chloroplast and human mitochondrion.**

A

**Ebola**

| TIS (real) | TIS (predict) | TTS (real) | TTS (predict) | ORF Name |
|---|---|---|---|---|
| 469 | 469 | 2686 | 2686 | NP |
| 3128 | 3128 | 4148 | 4148 | VP35 |
| 4478 | 4478 | 5456 | 5456 | VP40 |
| 6038 | 6038 | 8065 | 8065 | GP |
| | 7060 | | | |
| 8508 | 8508 | 9372 | 9372 | VP30 |
| 10344 | 10344 | 11097 | 11097 | VP24 |
| 11580 | 11580 | 18216 | 18216 | L |

B

**SARS-CoV-2**

| TIS (real) | TIS (predict) | TTS (real) | TTS (predict) | ORF Name |
|---|---|---|---|---|
| 265 | 265 | 13480 | | ORF1a |
| 265 | 265 | 21552 | 21552 | ORF1ab |
| | 1604 | | | |
| | 13767 | | | |
| 21562 | 21562 | 25381 | 25381 | S |
| 25392 | 25392 | 26217 | 26217 | ORF3a |
| | 25404 | | | |
| 26244 | | 26469 | | E |
| 26522 | 26522 | 27188 | 27188 | M |
| | 26771 | | 27266 | |
| 27201 | | 27384 | 27384 | ORF6 |
| 27393 | 27393 | 27756 | 27756 | ORF7a |
| 27755 | | 27884 | 27884 | ORF7b |
| 27893 | 27893 | 28256 | | ORF8 |
| 28273 | 28273 | 29530 | 29530 | N |
| | 28283 | | 29160 | |
| | 28573 | | 29292 | |
| | 28640 | | 29330 | |
| | 28900 | | 29334 | |
| | 28972 | | 29400 | |
| | 29221 | | 29447 | |
| | | | 29504 | |
| | | | 29514 | |
| 29558 | | 29671 | | ORF10 |
| | | | 29803 | |

**Figure S6. All predicted TISs and TTSs of Ebola and SARS-CoV-2.**
